# High-throughput identification of functional regulatory SNPs in systemic lupus erythematosus

**DOI:** 10.1101/2023.08.16.553538

**Authors:** Qiang Wang, Taehyeung Kim, Marta Martínez-Bonet, Vitor R. C. Aguiar, Sangwan Sim, Jing Cui, Jeffrey A. Sparks, Xiaoting Chen, Marc Todd, Brian Wauford, Miranda C. Marion, Carl D. Langefeld, Matthew T. Weirauch, Maria Gutierrez-Arcelus, Peter A. Nigrovic

**Author notes:** Co-first authors (Qiang Wang, Taehyeung Kim). **Correspondence:** Peter A. Nigrovic, MD Division of Immunology Boston Children’s Hospital Karp Family Research Building 10211 One Blackfan Circle Boston, MA 02115. **Conflict of interest:** The authors declare no relevant competing interests.

## Abstract

Genome-wide association studies implicate multiple loci in risk for systemic lupus erythematosus (SLE), but few contain exonic variants, rendering systematic identification of non-coding variants essential to decoding SLE genetics. We utilized SNP-seq and bioinformatic enrichment to interrogate 2180 single-nucleotide polymorphisms (SNPs) from 87 SLE risk loci for potential binding of transcription factors and related proteins from B cells. 52 SNPs that passed initial screening were tested by electrophoretic mobility shift and luciferase reporter assays. To validate the approach, we studied rs2297550 in detail, finding that the risk allele enhanced binding to the transcription factor Ikaros (IKZF1), thereby modulating expression of *IKBKE*. Correspondingly, primary cells from genotyped healthy donors bearing the risk allele expressed higher levels of the interferon / NF-κB regulator IKKε. Together, these findings define a set of likely functional non-coding lupus risk variants and identify a new regulatory pathway involving rs2297550, Ikaros, and IKKε implicated by human genetics in risk for SLE.

## Introduction

Systemic lupus erythematosus (SLE) is a chronic autoimmune disease predominantly affecting women. Identical twin concordance rates range between 30–50%, and first-degree relatives exhibit an approximately 20-fold increased risk of SLE compared with the general population [1–3]. While monogenic SLE has proven extremely informative with respect to causative pathways, most SLE is polygenic, arising through a complex interplay of genetic and environmental factors [4, 5]. Correspondingly, genome-wide association studies (GWAS) have defined dozens of loci that modulate disease risk [6–10]. Deciphering these different genetic contributors, each individually minor but collectively of major importance, will be central to defining SLE heterogeneity at a genetic level.

Unfortunately, translating GWAS data into pathogenic understanding has proven difficult. Few common disease-associated variants affect coding regions, indicating that the associated variants – predominantly single-nucleotide polymorphisms (SNPs) – likely exercise a regulatory role. Distinguishing functional non-coding SNPs from the many incidental SNPs that reside in close linkage disequilibrium (LD) remains a “needle in the haystack” problem of great complexity. Strategies to address this problem in polygenic diseases include fine-mapping [11], proteome-wide profiling of DNA-binding proteins over SNPs [12], integration of expression quantitative trait loci (eQTL) with epigenetic and/or transcriptomic data [13–15], massively parallel reporter assays [16–18], and SNP-seq [19].

Here, we pursue a combined experimental and bioinformatic approach to identify functional non-coding variants in SLE that act by regulating the binding of transcription factors and other DNA-binding proteins. We focus on B cells as a lineage implicated genetically as critical to SLE pathogenesis [20]. We used SNP-seq to screen for alleles that confer differential binding to B cell nuclear proteins, enriching targets further through epigenetic markers of open chromatin and predictors of regulatory function. Beginning with 2180 candidates, we identify 52 plausible regulatory SNPs, each interrogated experimentally to prioritize 5 high-likelihood functional variants. To test our approach, we examined rs2297550 in detail, finding that this variant modulates the binding of the transcription factor Ikaros to regulate expression of the noncanonical IκB kinase IKKε. Thus, our strategy bridges the gap between GWAS and mechanism while defining a novel regulatory pathway implicated by population genetics in risk for SLE.

## Results

### Candidate functional non-coding variant identification via SNP-seq and bioinformatic screens

To identify functional non-coding variants associated with SLE, we began from GWAS hits with genome-wide significance (P value < 5 x 10^-8^) from five studies [7, 9, 10, 21, 22], finding 87 loci of interest, each defined by a tagging SNP (**Supplementary Table 1**). We identified all common variants in LD with these SNPs from 1000 Genomes Phase 3 (European population), filtering on minor allele frequency [MAF] >1%, LD R^2^>0.8, and location within 1 Mb of the tagging SNP (**Figure 1A**). This strategy yielded 2180 candidate SNPs for downstream screening and validation (**Supplementary Table 1**).

**Figure 1.**
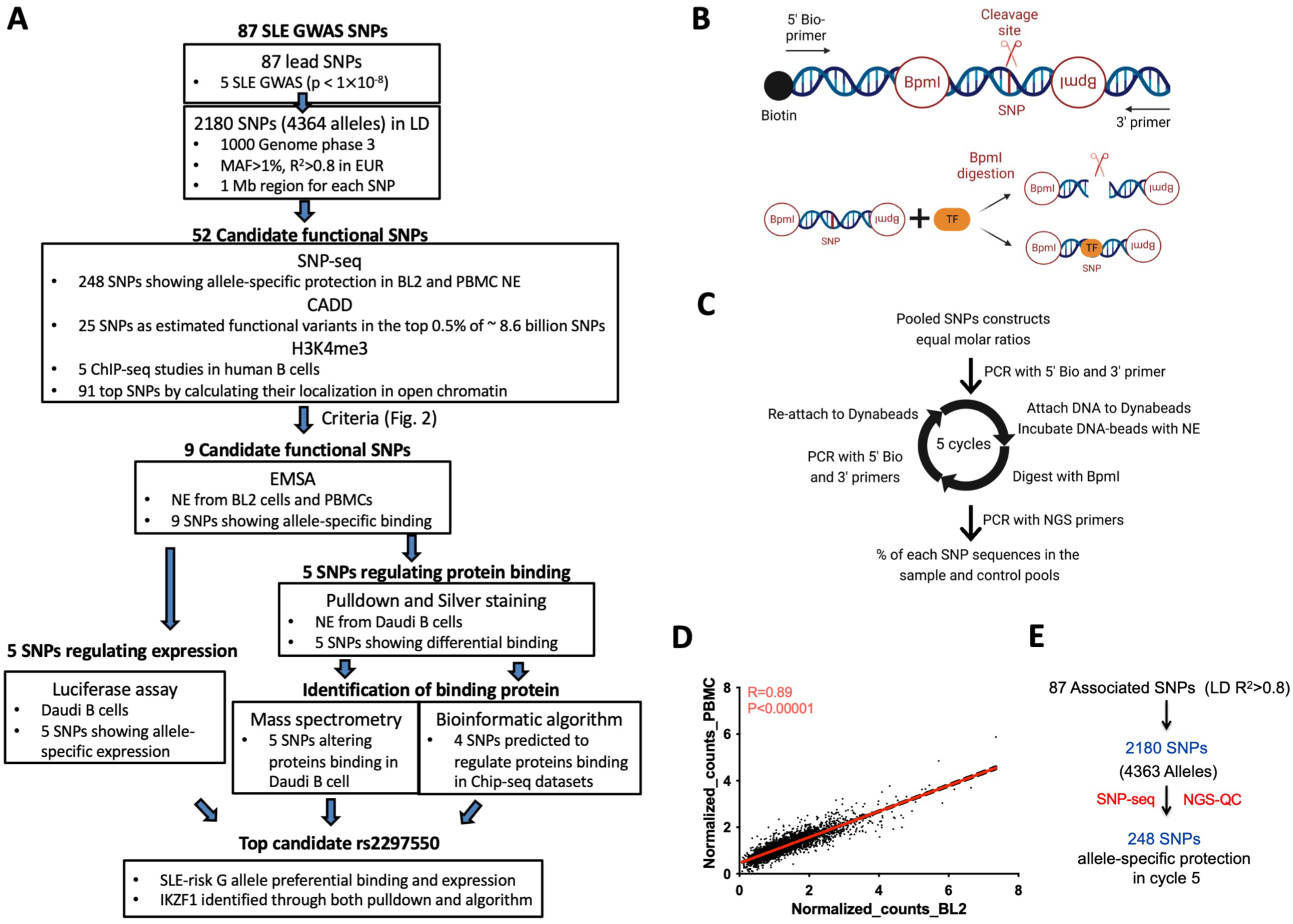
Screening candidate regulatory variants for SLE by SNP-seq. **(A)** Flow chart illustrating identification of functional candidate SNPs from SLE GWAS data. **(B)** SNP-seq: To generate the SNP-seq construct (top), a 31 bp sequence centered on the SNP is positioned between two type IIS restriction enzyme (IIS RE) binding sites. SNPs that fail to bind regulatory proteins such as transcription factors (TF) are negatively selected (bottom), allowing enrichment of protected constructs by PCR. The whole construct can be amplified using primers as per **Suppl. Table 8**. Bio, biotin. Created with BioRender. **(C)** The experimental procedure for SNP-seq; NE, nuclear extract; Bio, biotin. **(D)** Spearman’s correlation analysis of SNP-seq allele counts normalized to control between PBMC and BL2 samples. **(E)** We selected 248 SNPs that passed next generation sequencing quality control (NGS-QC) and demonstrated progressive allele-specific protection (**Suppl. Figure. 1C**).

To narrow this pool, we applied SNP-seq [19]. In this method, a library of double-stranded DNA constructs is generated that includes each allele of each candidate SNP. A 31bp sequence centered on each SNP is flanked by binding sites for the type IIS restriction enzyme BpmI that cleaves DNA in a sequence-independent manner 16bp (5’)/14bp (3’) adjacent to its binding site. Binding of transcription factors or other nuclear proteins to the SNP site protects the construct from enzymatic digestion (**Figure 1B**), as recognized by subsequent amplification and DNA sequencing [19, 23]. Here, our library contained 4363 dsDNA constructs reflecting each allele of the 2180 SNPs, including 3 with 3 alleles (**Supplementary Table 2**). The library was then incubated with nuclear extract from either human peripheral blood mononuclear cells (PBMCs) from healthy individuals or BL2 B cells, washed, and then digested with BpmI. As a control, the construct pool was incubated without nuclear extract. The remaining sequences were amplified by PCR and used as the library for the next round of selection. The whole procedure was repeated for 5 cycles, with barcoding at cycles 3 and 5 to identify enriched SNPs by next-generation sequencing (NGS) (**Figure 1C**). The entire procedure was performed in three independent biological replicates, finding high correlation (Spearman ρ>0.8) between sample pairs (**Supplementary Figure 1A, 1B**) and between PBMC and BL2 samples (**Figure 1D**).

Two analytical approaches were employed to identify SNPs offering allele-specific protection, as established previously [19]. First, we chose SNPs that exhibited at least 20% difference in protection between alleles in cycle 5 in samples treated with nuclear extract from both PBMC and BL2 cells; 452 SNPs exhibited such an allelic differential. Second, we sought SNPs demonstrating progressive allele-specific protection across cycles (cycle 0, 3, and 5) (**Supplementary Figure 1C**); 496 SNPs exhibited this pattern. In the end, 248 SNPs met both criteria (**Figure 1E, Supplementary Table 3**). A diagram of our analytical strategy is shown in **Supplementary Figure 2**.

Next, we examined the same 2180 SNPs using two bioinformatic approaches. First, we employed informative chromatin marks to identify areas of open chromatin, calculating H3K4me3 chromatin scores from 5 chromatin immunoprecipitation (ChIP)-seq studies in human B cells in the Encyclopedia of DNA Elements (ENCODE) database, as per a described method [24] (**Supplementary Figure 3**). We selected for further study those SNPs that ranked in the top 100 from at least half of studies, yielding 91 variants (**Supplementary Table 4**). Second, we used Combined Annotation-Dependent Depletion (CADD), a composite measure that employs 60 genomic features to estimate the probability that a variant is functional [25]. We ranked all 2180 SNPs as per the CADD database (https://cadd.gs.washington.edu/), retaining the 25 SNPs that scored in the top 0.5% of all ∼8.6 billion single nucleotide variants (**Supplementary Table 5**).

To reduce the number of SNPs to a tractable number for experimental validation, we chose the top 20 SNPs identified uniquely by SNP-seq, the top 8 SNPs identified uniquely by epigenetic and CADD analysis, and 18 SNPs identified in at least two of three analyses (6 SNP-seq+H3K4me3, 4 SNP-seq+CADD, 8 H3K4me3+CADD), for a total of 52 plausible functional SNPs (**Figure 2**, **Supplementary Table 6**).

**Figure 2.**
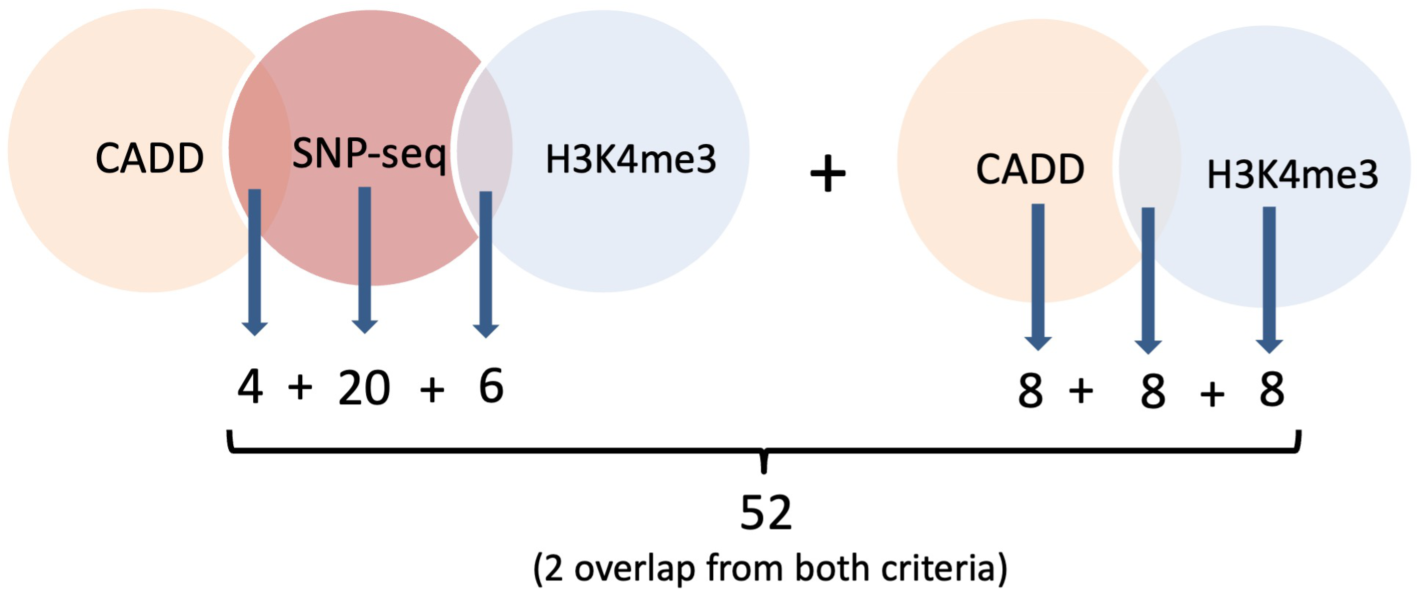
Prioritization of candidate SNPs by SNP-seq and bioinformatic enrichment. SNPs selected for experimental screening were the top 20 of 248 emerging form SNP-seq, the top 8 emerging from H3K4me3 score, the top 8 emerging from CADD, and 18 SNPs identified by at least two analyses: 6 SNP-seq+H3K4me3, 4 SNP-seq+CADD, and 8 SNPs H3K4me3+CADD.

### Experimental validation of plausible functional SNPs in SLE

As experimental validation of the 52-SNP enriched variant pool, we first employed the electrophoretic mobility shift assay (EMSA). Biotinylated 31bp DNA fragments centered on each allele of each the 52 candidate SNPs (111 alleles) were interrogated using nuclear extract from BL2 cells (triplicate biological repeats) followed by a validation EMSA using nuclear extract from human PBMCs. Overall, 7 biallelic and 2 triallelic (rs2297550 and rs936394) SNPs exhibited allele-imbalanced binding for all nuclear extracts: rs2297550, rs906868, rs936394, rs7302634, rs276461, rs13213604, rs378056, rs2814955 and rs9907966

(**Figure 3A**, shift binding bands marked as red circles). All EMSA blots for every experiment and a summary table can be found in **Supplementary Figures 4 and 5**.

**Figure 3.**
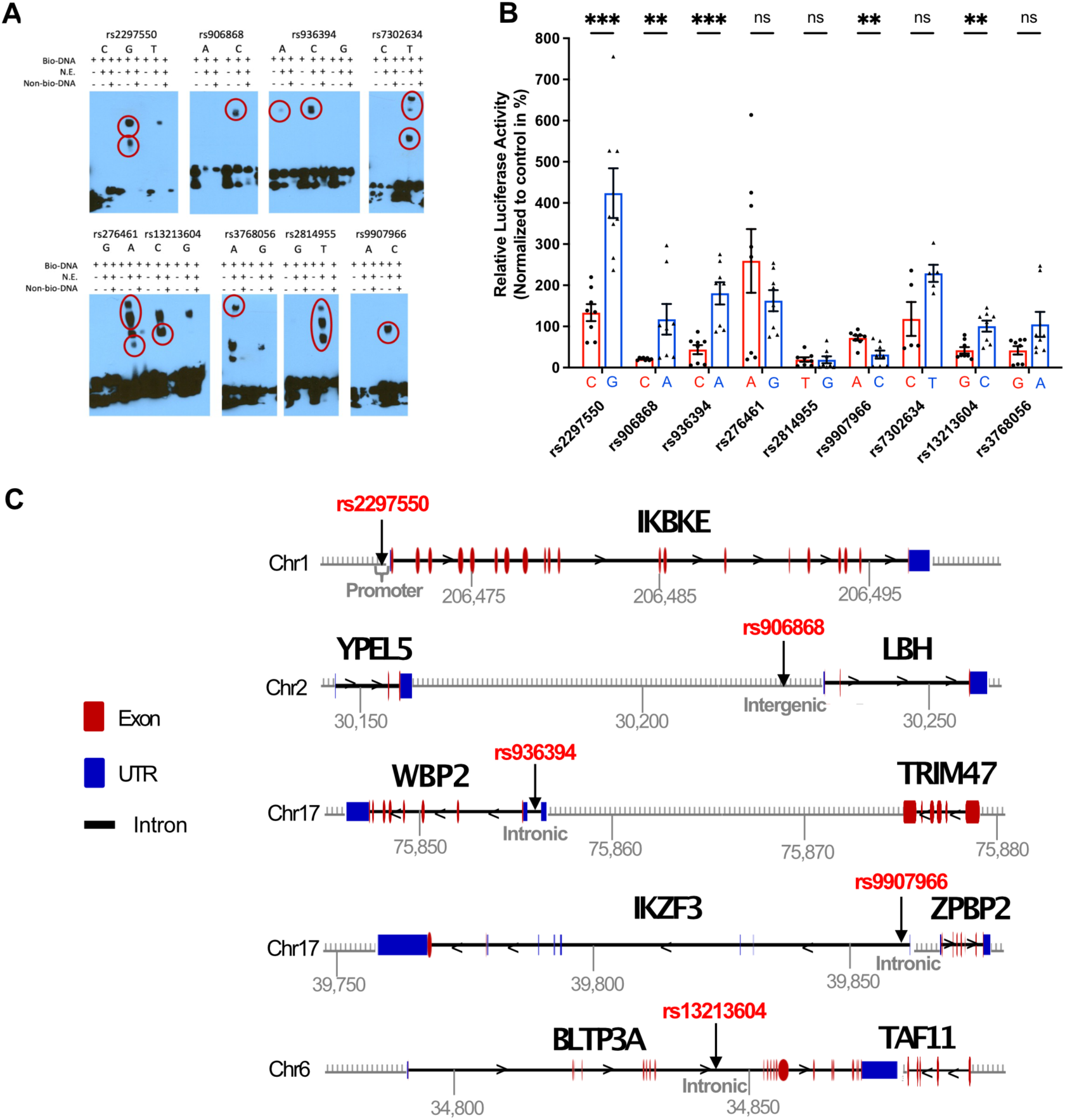
EMSA and luciferase reporter assessment of 52 candidate regulatory SNPs. **(A)** 7 biallelic and 2 triallelic (rs2297550 and rs936394) SNPs showed consistent allele-imbalanced binding for nuclear by EMSA. Allele-specific gel shift/binding marked with red circles. PBMC data shown; BL2 data and negative SNPs provided in **Suppl. Fig.4**. **(B)** Luciferase reporter assay between the reference (red) and alternative (blue) alleles of 9 candidate regulatory SNPs from A in Daudi B cells (mean ± s.d, n=8 biological replicates, Mann-Whitney test with two-stage step-up correction for multiple hypothesis testing: rs2297550 p=0.00047, rs906868 p=0.0013, rs936394 p= 0.00047, rs9907966 p=0.0071, rs13213604 p=0.0085). **(C)** Diagram displaying the position of 5 SNPs that showed consistent significant differences between alleles through EMSA and luciferase reporter assay; rs2297550 in promoter of *IKBKE*, rs906868 in intergenic region between *YPEL5* and *LBH*, rs936394 in intron of *WBP2*, rs9907966 in intron of *IKZF3*, rs13213604 in intron of *BLTP3A*. The unit of chromosome position is in kilobases (Kb). Diagram of gene features including exon, intron, and UTR was generated using GSDS 2.0 (http://gsds.gao-lab.org/index.php).

Next, we assessed the allele-specific gene regulatory capacity of these 9 SNPs using a luciferase reporter assay; for rs2297550 and rs936394, we included only two alleles, since the third for each has a MAF <0.1% and is thus unlikely to mediate GWAS signal. We cloned the same risk or non-risk allele DNA fragments as for EMSA (but without biotinylation) into the pGL3 vector, and then transfected each construct into Daudi B cells, simpler to transfect than BL2, together with control vector pRL to normalize transfection efficiency. We found 5 SNPs to retain a significant difference between alleles and thus represent high-probability functional non-coding variants: rs2297550, rs906868, rs936394, rs9907966 and rs13213604 (**Figure 3B**). These SNPs are diagrammed in **Figure 3C**.

### Identification of potential transcription factor binding

To provide independent validation of the identified SNPs, we sought to confirm binding of transcription factors and/or other nuclear proteins in an allele-determined manner. Again, we approached this problem both experimentally and bioinformatically.

First, we employed pulldown and silver staining in SNPs that had exhibited allele-specific differential binding by EMSA, excluding rs2814955 because of its low luciferase activity, again with 31bp oligonucleotides for each allele but now using nuclear extract from Daudi cells [26]. In two biological repeats, four high-probability functional variants again showed differential binding (red circles) (**Supplementary Figure 6**). Each pulldown product for each allele of the five high-probability SNPs was evaluated by mass spectrometry to identify binding nuclear proteins, yielding a set of candidate proteins for each (**Supplementary Table 7**). Second, we employed a bioinformatic algorithm based on published ChIP-seq data to identify candidate transcription factors likely to recognize each of the 5 high-probability variants [27, 28]. Bioinformatic predictions are shown in **Supplementary Table 8**. Evaluating these data together, we observed that the transcription factor Ikaros (IKZF1) was identified in the pulldown product for rs2297550 and was predicted by the algorithm, including preferential binding to the SLE-risk G allele as observed by EMSA. We therefore chose rs2297550 as a test case for our selection process.

### rs2297550 regulates *IKBKE*/IKKε expression

As depicted in **Figure 3C**, rs2297550 resides in a promoter/enhancer region, 47bp upstream of *IKBKE*, encoding the kinase IKKε [29–31]. This SNP emerged from Chinese and European GWAS data as a leading candidate in SLE, with *IKBKE* as its likely target gene [7, 32]. To assess the role rs2297550 in regulation of *IKBKE*, we used CRISPR homology directed repair to generate base-edited Daudi cell clones with CC, CG, and GG genotypes. GG clones exhibited lower mRNA for *IKBKE* (**Figure 4A**) and lower expression of IKKε protein (**Figure 4B**), confirming a regulatory relationship between SNP and gene. Comparable findings were observed in LPS-stimulated Daudi cells (**Supplementary Figure 7**).

**Figure 4.**
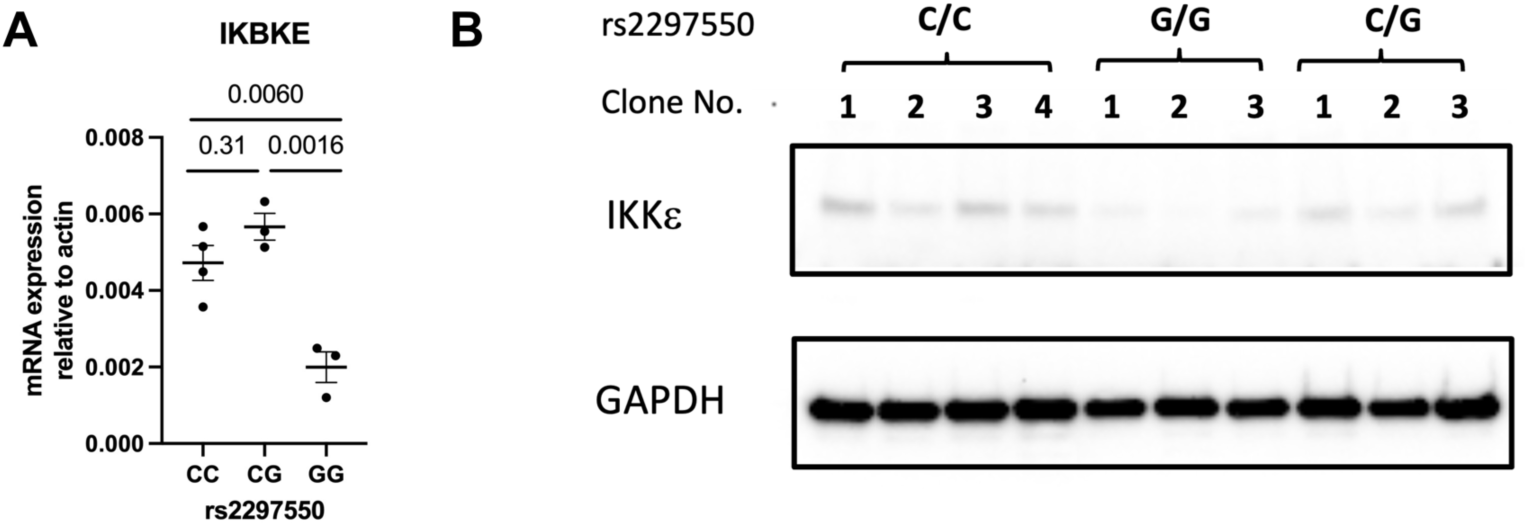
Characterization of functional SNP rs2297550 by CRISPR-mediated editing in Daudi cells. **(A)** mRNA levels of *IKBKE* and 𝛽-actin by RT-qPCR and **(B)** protein levels of IKK𝜖 (80 KDa) and GAPDH (37 KDa) were measured from base-edited 4 C/C, 3 C/G, and 3 G/G clones (mean ± s.d, *P* values from one-way ANOVA corrected by Tukey’s multiple comparisons).

Next, we recruited healthy donors bearing different genotype of rs2297550 to test the impact of genotype in primary PBMCs, using the Joint Biology Consortium (JBC) patient recruitment infrastructure and the Mass General Brigham Biobank (www.jbcwebportal.org). In resting cells, IKKε protein levels were comparable across genotypes in PBMCs collectively and in all immune subtypes tested (CD19+ B cells, CD3+CD4+ T cells, CD3+CD8+ T cells, CD3-CD56+ NK cells, CD14+ monocytes) (**Figure 5A-F**).

**Figure 5.**
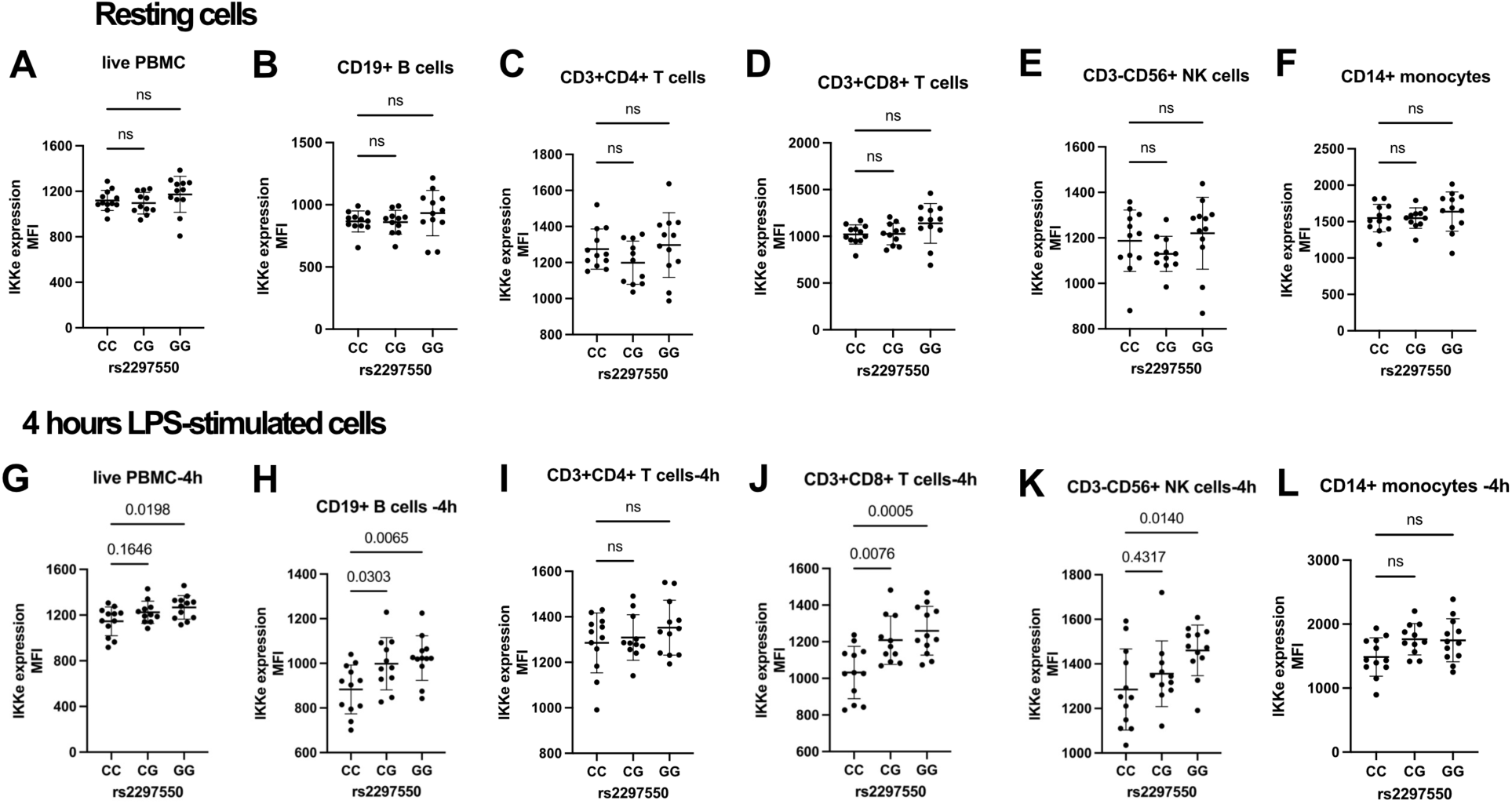
SNP rs2297550 associated with IKK𝝐 expression in primary PBMCs. IKK𝜖 expression was determined by flow cytometry for 35 healthy subjects with C/C (n=12), C/G (n=11), G/G (n=12) genotype at SNP rs2297550. The mean fluorescence intensity (MFI) of IKK𝜖 expression in (**A-F**) resting and (**G-L**) 4 hours LPS-stimulated PBMCs, including CD19+ B cells, CD3+CD4+ T cells, CD3+CD8+ T cells, CD3-CD56+ NK cells, and CD14+ monocytes (mean ± s.d, *P* values from one-way ANOVA corrected by Dunnett’s multiple comparisons test).

However, after 4h of LPS stimulation, cells from GG donors exhibited differential protein levels in PBMCs generally as well as in B cells, CD8+ T cells, and NK cells; a heterozygote (allele dose) effect was observed in B cells and CD8+ T cells (**Figure 5G-L**). Interestingly, unlike in Daudi cells, the effect of the risk allele was to increase rather than decrease IKKε expression.

### Ikaros binds to rs2297550 to regulate *IKBKE*

Given concordant findings from mass spectrometry and bioinformatic prediction, we pursued the possibility that Ikaros binds to rs2297550 to regulate *IKBKE*. We therefore performed an oligonucleotide pulldown assay using nuclear extract from Daudi cells, with or without excess non-biotinylated competitor, followed by western blot using anti-Ikaros antibody. Ikaros bound both C and G alleles but exhibited a preference for the G allele, as predicted, reflected in the disappearance of signal with excess non-biotinylated G competitor (**Figure 6A**). To confirm this result, we employed EMSA-supershift, finding that anti-Ikaros weakened binding to rs2297550-G compared with isotype control or antibody to another transcription factor candidate from the bioinformatic prediction, E2F transcription factor 4 (E2F4) (**Figure 6B**). These studies establish that Ikaros binds rs2297550 and that this interaction is allele specific.

**Figure 6.**
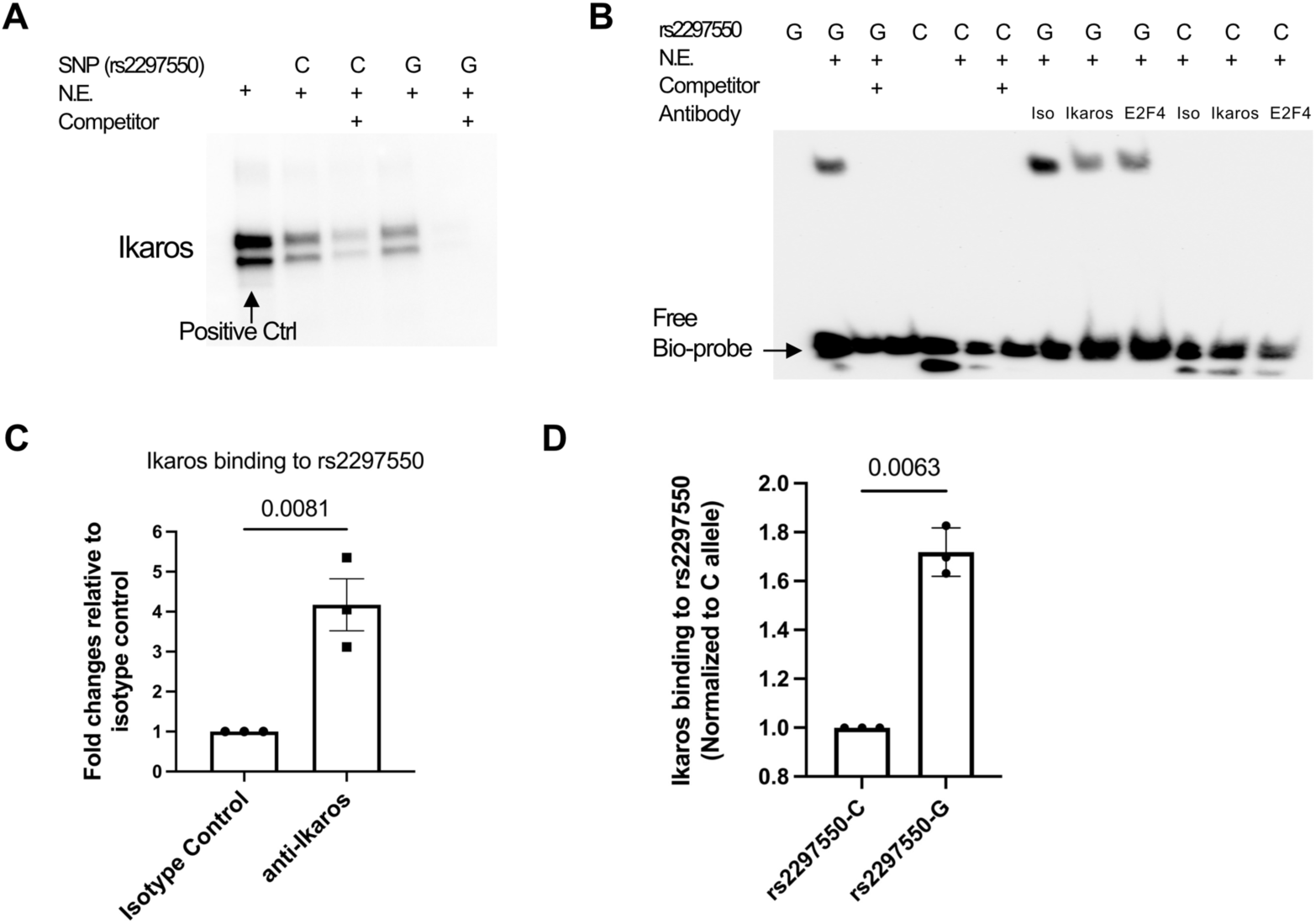
rs2297550 risk allele G bind Ikaros. **(A)** Oligonucleotide pulldown western blot assay testing Ikaros binding to rs2297550-G; binding is eliminated in the presence of a 30-fold excess of non-biotinylated rs2297550-C or G competitor; positive control, nuclear extract. **(B)** EMSA supershift assay showing that anti-Ikaros caused fading of the shifted band (oligo rs2297550-G, Daudi nuclear extract), whereas isotype control did not; anti-E2F4 also exhibited specificity. **(C)** ChIP-qPCR using anti-Ikaros confirms binding to rs2297550 (Daudi cells, mean ± s.d, n=3). (**D**) Allele discrimination ChIP-qPCR shows Ikaros preferentially binds to G allele over C allele of rs2297550 in CRISPR-HDR edited C/G Daudi cells (mean ± s.d, n=3). All studies done in unstimulated cells. Statistical analysis for C and D: paired *t* test with two tails, without correction.

To assess rs2297550-Ikaros binding in living cells, we performed chromatin immunoprecipitation (ChIP)-qPCR. In Daudi cells, anti-Ikaros antibody precipitated rs2297550 significantly more than did isotype control antibody (**Figure 6C**). Further, in CRISPR-HDR edited CG Daudi cells, we observed preferential pulldown of the G allele over the C allele (**Figure 6D**), again confirming its allelic preference.

To test whether the interaction of Ikaros with rs2297550 modulated IKKε expression, we generated three Ikaros knockout clones using CRISPR-Cas9. All clones showed Ikaros deficiency (**Figure 7A**) and corresponding loss of the ability of anti-Ikaros to pull down rs2297550 (**Figure 7B**). Consistent with the inhibitory effect of G allele carriage in these cells, loss of Ikaros increased *IKBKE* mRNA (**Figure 7C**) and IKKε protein (**Figure 7D**). These findings establish a new rs2297550-Ikaros-IKKε regulatory axis implicated by human population genetics in risk for SLE.

**Figure 7.**
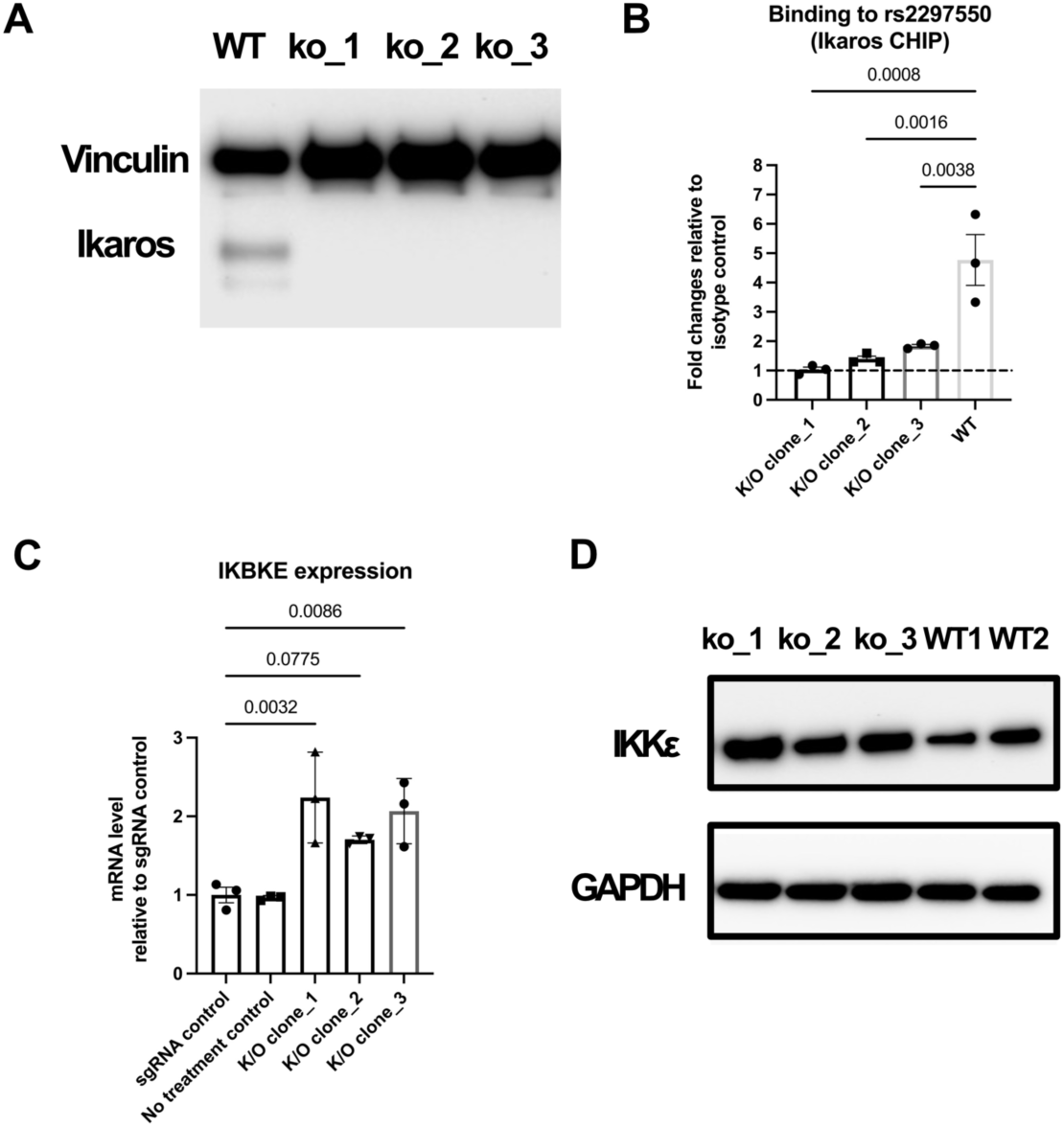
Ikaros binding to rs2297550 regulates IKBKE/IKK𝝐 expression. CRISPR-Cas9 Ikaros knockout Daudi clones show **(A)** decreased Ikaros protein expression by western blotting, **(B)** loss of Ikaros binding to rs2297550 by ChIP-qPCR (mean ± s.d, n=3), and **(C, D)** increased *IKBKE* mRNA by RT-qPCR (mean ± s.d, n=3) and increased IKK𝜖 protein expression by western blotting. Statistical analysis panels B and C: one-way ANOVA corrected by Dunnett’s multiple comparisons test.

**Figure 8.**
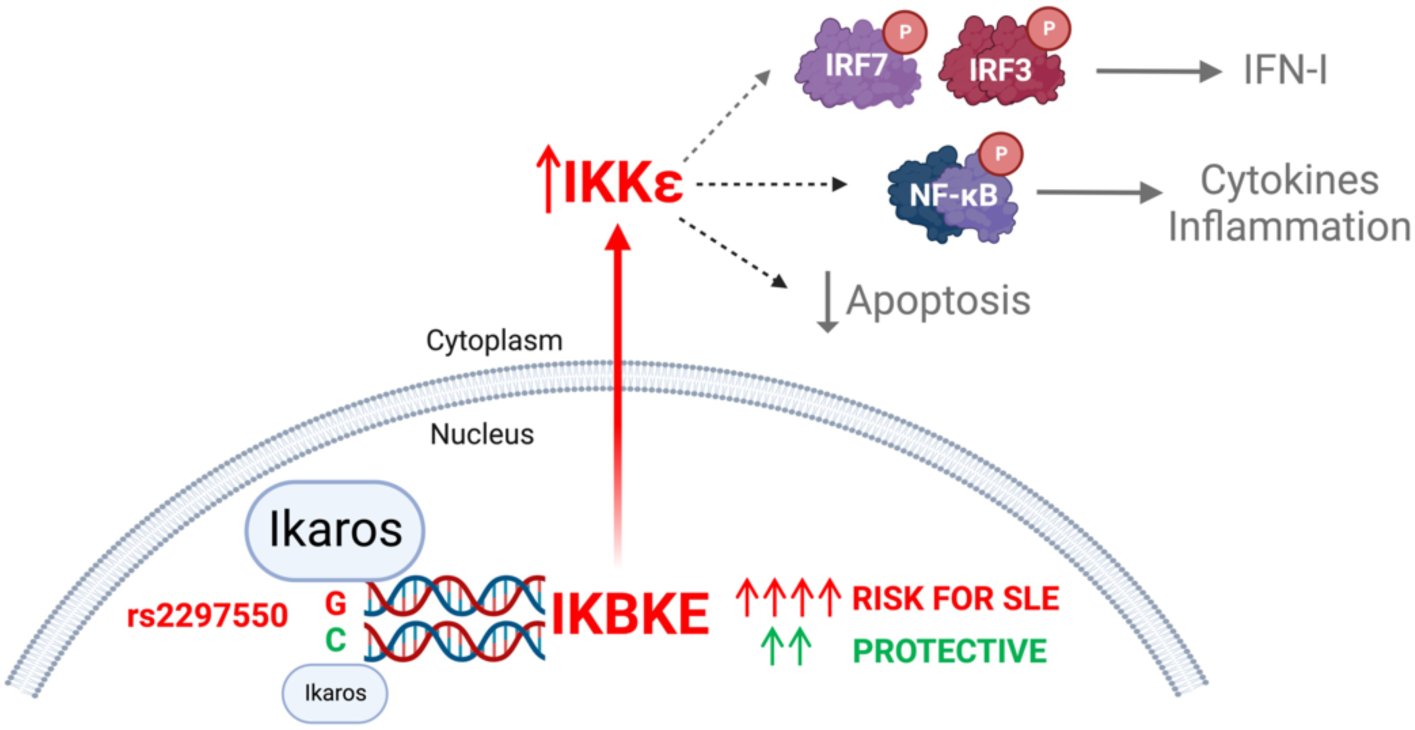
The lupus risk SNP rs2297550 allelically modulates binding of the transcription factor Ikaros (encoded by *IKZF1*) to regulate expression of IKBKE/IKKε, a protein that modulates multiple processes implicated in SLE pathogenesis. Note that Ikaros binding can either amplify or repress IKBKE depending on context; amplification, shown here, reflects the direction of effect observed in genotyped healthy donors. The role of mechanisms downstream of IKKε remains to be defined. Created with BioRender.

## Discussion

Genetic risk for SLE is borne by many susceptibility loci, most of which reflect common non-coding variants [33]. Despite the modest effect of particular variants on disease risk, each unambiguously implicates its associated pathway in the pathogenesis of human disease. Further, as the number of risk variants accumulates, incidence of SLE increases and age of onset declines, suggesting a “cumulative hit” model whereby variants operate synergistically to generate disease susceptibility [9]. Understanding which specific variants drive risk and how they do so offers a unique window into disease biology as well as into general mechanisms of immune control [34, 35].

Here, we conducted a systematic experimental and bioinformatic screen across 87 loci implicated by GWAS in SLE, identifying 52 plausible casual non-coding SNPs, of which 5 emerged through EMSA and luciferase as high-likelihood variants. Extrapolation from surrounding genes suggests that these 5 variants likely interface with functional pathways relevant for SLE biology, including the interferon pathway (*IKBKE*), pyroptosis (*IKBKE*), B cell differentiation (*IKZF3*) and WNT signaling (*LBH*) (**Figure 3C**).

To test our approach, we studied one variant in detail: rs2297550, a triallelic SNP (frequency of the three alleles C 0.862, G 0.138, T <0.001 per gnomAD) that resides 4462 bp upstream of the start codon for *IKBKE*, encoding the kinase IKKε. The minor allele G confers an increased risk of SLE with an odds ratio of 1.16 (95% CI: 1.11-1.21, p=1.31 × 10^−11^) compared with the major C allele [7]. Base editing showed that allelic variation at rs2297550 changes expression of *IKBKE* / IKKε. Variation in IKKε with genotype at rs2297550 was confirmed in healthy volunteer donors. As an orthogonal line of evidence, we showed that Ikaros exhibited allele-dependent affinity for rs2297550 and that deletion of Ikaros altered *IKBKE*/IKKε expression. These results establish a previously undefined rs2297550-Ikaros-IKKε regulatory axis.

Some groups have found that the risk allele G at rs2297550 is associated with lower *IKBKE* mRNA in unstimulated PBMCs [31, 36]. IKKε protein was not tested, raising the possibility that transcript and protein levels could be discordant. Indeed, we observed a trend toward lower levels of *IKBKE* mRNA in GG PBMC cells, though this effect did not reach significance (**Supplementary Figure 8**). The basis for transcript-protein divergence is not clear but could relate to the relative half-lives of mRNA and protein, among other factors [37, 38].

Interestingly, the relationship between genotype and IKKε varied with context. In Daudi B lymphoblasts, a line derived from a patient with Burkitt lymphoma, introduction of the risk GG genotype decreased IKKε expression and deletion of Ikaros enhanced it, establishing Ikaros as an inhibitor of *IKBKE*. This effect persisted after stimulation with LPS. By contrast, in healthy donors, circulating B cells showed an *increase* in IKKε with risk allele carriage, including a gene dose effect, but only after LPS stimulation.

Similar regulation was observed in CD8+ T cells, NK cells, and monocytes. The dependence of the impact of genetic variants on cell type and developmental/activation state is well recognized [39]. Since Ikaros can toggle between activator and repressor depending on associated cofactors, it may be that the variation observed here reflects differential expression of these proteins, or on other genetic/epigenetic differences between the lineages studied [40]. Under conditions other than those tested here, it is possible that the risk allele at rs2297550 could suppress IKKε in primary cells, as it does in Daudi cells.

An important intracellular intermediate in innate immune responses, IKKε can activate IRF3, IRF7, and NF-κB pathways to drive gene transcription in response to extracellular and intracellular stimuli, resulting in production of inflammatory mediators including type I interferons (**Figure 8**) [41–44]. The known role of interferons in SLE makes this pathway an appealing potential explanation for why rs2297550 is a risk SNP for SLE [45]. IKKε can also protect cells from TNF-induced apoptosis, plausibly thereby attenuating autoantibody production [46, 47]. *IKBKE* mRNA expression has been reported to be either higher [48, 49] or lower [31, 50, 51] in whole-blood and/or PBMC of SLE patients compared with healthy donors. Our primary cell data suggest that the risk phenotype attributable to rs2297550 likely reflects enhanced rather than suppressed expression of IKKε, though further work will be required to define the mechanistic connection between IKKε and SLE.

*IKZF1* encodes Ikaros, an essential regulator of lymphocyte development [40, 52]. Ikaros is implicated in SLE by multiple lines of evidence. A region including *IKZF1* carries genetic risk for SLE [32, 53–57]. Individuals bearing mutant *IKZF1* can present with SLE-like autoimmune disease [58, 59]. Clinical efficacy has been established in human SLE for iberdomide, an agent that accelerates degradation of Ikaros as well of the related transcription factor Aiolos [60, 61]. Mechanistically, Ikaros is involved in B cell tolerance and plasmacytoid dendritic cell differentiation [40, 62–64]. In B cells from SLE patients, iberdomide attenuated in vitro differentiation to plasmablasts and plasma cells as well as antibody production [65]. Thus, our findings implicating Ikaros in SLE risk via rs2297550 are consistent with genetic, clinical, and mechanistic observations in SLE.

Our study establishes a new connection between Ikaros, IKKε, and SLE, showing that the regulation of this intracellular signaling molecule is part of the portfolio of this transcription factor, and implicating both proteins in risk of incident SLE in humans. This finding has potential therapeutic relevance, since drug candidates supported by human genetic evidence are much more likely to succeed in clinical trials than those without such support [66, 67]. However, our data illustrate the challenge of extrapolating from a genetic variant to a single mechanism. Depending on cellular context, we find that the same variant has opposite effects on gene expression. Further, while we began our search in B cells, in human primary cells the rs2297550-IKKε effect spans multiple immune lineages, both adaptive and innate. Thus our data do not establish which lineages, and which pathways downstream of IKKε, contribute to the mechanisms through which rs2297550 drives SLE risk.

Other limitations arise from our methodology. Non-coding functional variants may operate in ways not detectable by SNP-seq or by our bioinformatic screens, for example through chromosome conformation, microRNA, and mRNA splicing. We would not have detected variants that operate via DNA-binding proteins absent from the nuclear extract employed, for example, transcription factors specific for other B cell developmental/activation states or in different immune or non-immune lineages. For this reason, while the high-probability SNPs described here are likely functional, we expect that many genuinely functional non-coding SNPs will have been filtered out. Definitive validation of the remaining 4 high-probability SNPs will require dedicated study of the kind devoted here to rs2297550. While our findings suggest the possibility of epistasis between *IKBKE* and *IKZF1*, inadequate resolution of the IKZF1 locus in the Immunochip rendered an attempt to identify such an interaction impossible [9]. Thus, the existence of an epistatic interaction between these loci remains to be tested.

Our studies in genotyped donors employed healthy donors rather than SLE patients. This choice was deliberate. We derived our initial candidate SNPs from GWAS that compared patients with SLE to patients lacking SLE. This design identifies variants modulating the risk that a healthy person will develop disease, and therefore that must be operative before SLE appears – that is, in healthy individuals. Such variants may also modulate disease phenotype, severity, or response to therapy in patients with established SLE, but such effects remain conceptually distinct from how they predispose to incident disease, as studied here.

Our combined experimental and bioinformatic approach also has clear strengths. The number of SNPs that survived each individual filter was substantial, illustrating the difficulty of separating functional from incidental SNPs. Multiple orthogonal approaches are required to validate any individual SNP and remains highly labor-intensive. The value of such an effort arises from the fact that each functional variant points the way to a pathway relevant in human SLE. It is key to remember that the effect size of a non-coding variant reflects the impact of the polymorphism on disease risk, but not the importance of the gene itself in disease pathogenesis, since regulatory SNPs often alter gene expression only modestly, exactly as observed here. Further, non-coding variants can be expected to identify pathways that coding variants cannot, either because mutations that disrupt a gene are poorly tolerated and thus eliminated from the population or because they lead to a phenotype that is different from that induced by a more subtle alteration in gene expression. As a result, any effort to understand highly polygenic disease must seek to understand not only highly penetrant coding variants but also each non-coding variant.

In summary, we provide a combined experimental and bioinformatic approach to identify plausible regulatory non-coding variants in SLE, proceeding from our initial screen to establish a previously unrecognized pathway through which Ikaros regulates IKKε, as modified by common allelic variation at rs2297550. Further investigation is required to understand how IKKε participates in SLE, including which lineages and downstream pathways are essential to this role. Our approach is broadly applicable and will help to translate GWAS into the pathogenic understanding that is a prerequisite to the personalized treatment of polygenic diseases such as SLE.

## Methods

### Cells and culture

Human B cell line BL2 was from DSMZ and human B cell line Daudi was kindly provided by Dr. Baglaenko Yuriy. BL2 and Daudi cells were cultured in RPMI1640 medium (ThermoFisher Scientific, catalog number 11875093) supplemented with 10% Fetal Bovine Serum (FBS) (ThermoFisher Scientific, catalog number 26140079) and 1% Penicillin-Streptomycin (ThermoFisher Scientific, catalog number 15140122).

### SNP-seq oligos and primers

All primers were purchased from IDT as listed in Supplementary Table 1. SNP-seq oligo pool (Supplementary Table 2) was purchased from TwistBiosciences and oligos for EMSA screening and luciferase reporter assay was purchased from EtonBiosciences.

### SNP-seq

SNP-seq constructs were built according to **Figure 1B**. The SNP sequence is 31 bp long DNA centered on the SNP of interest. The sequences of PCR amplification primers for the library oligonucleotides are listed in **Supplementary Table 9** and the library itself is detailed in **Supplementary Table 2**. For oligo library amplification, 100 ng of pooled DNA was amplified with bioMagF-G5 and MagR-G3 by PCR for 25 cycles with AccuPrime Taq (Thermo Fisher Scientific, catalog number 12346086) at 94 °C for 60 s, 58 °C for 60 s and 68 °C for 40 s. After gel purification, 10 ng of biotinylated DNA was attached to 4 μl streptavidin-Dynabeads (Invitrogen, catalog number 11205D) according to the manufacturer’s protocol. The DNA-bound beads were then incubated with 60 μg nuclear extract from BL2 cell line or PBMC or no nuclear extract for 1 h at room temperature in LightShift Chemiluminescent EMSA Kit reaction buffer (Thermo Fisher Scientific, catalog number 20148X). After washing and separation, the DNA-bound beads were digested with 2 μl BpmI (NEB, catalog number R0565L) for 30 min at 37 °C. After another wash and separation, the DNA was amplified again with bioMagF and MagR and reattached to the Dynabeads for the next SNP-seq cycle. Five cycles were performed in total. DNA from cycle 0, 3, 5 were used to prepare a Next Generation Sequencing (NGS) library by two consecutive PCRs (PCR with L1seq and R1 primers, 20 cycles of 98 °C 1:30 min, 60 °C 1:40 min, 72 °C 0:40 min; PCR with L2 and R2 primers, 25 cycles of 98 °C 1:30 min, 60 °C 1:40 min, 72 °C 0:40 min) for each independent sample using Herculase polymerase (Agilent, catalog number 600677). The primers used are also listed in **Supplementary Table 9.** PCR products were run in a 2% agarose gel and the correct band (around 200 bp) was purified with QiaQuick gel extraction Kit (QIAGEN, catalog number 28706) (at least 2 columns) following manufactureŕs instructions. The elution was sent for NGS at the Dana Farber Genomics Core, Harvard Medical School.

### Human PBMC isolation

Human peripheral blood mononuclear cells (PBMCs) used in SNP-seq were extracted from apheresis leukoreduction collar blood via Ficoll gradient (Cytiva, catalog number GE17-1440-02). For studies in genotyped donors, whole blood was collected from genotyped healthy human subjects recruited from the Mass General Brigham Biobank through Joint Biology Consortium (JBC) recruitment core (www.jbcwebportal.org). Genotype was confirmed in each donor by Sanger sequencing. PBMCs were isolated from whole blood collected in BD Vacutainer™ Plastic Blood Collection Tubes with K2EDTA via Ficoll gradient and stored in liquid nitrogen in 10% DMSO in complete FBS.

Human subjects research was approved under Brigham and Women’s Hospital IRB protocol 2019P003709 and the Boston Children’s Hospital IRB protocol P000043108.

### Chromatin marks and CADD

Chromatin mark score calculation is based on published protocol [24]. In short, for a variant, the score is the ratio of H3K4me3 epigenetic peak value from CHIP-seq data to the distance between variant and its closest H3K4me3 peak. Five H3K4me3 CHIP-seq experiments for B cells from ENCODE are used for chromatin mark score calculation. The experimental ID and final sheet for all the scores are provided in **Supplementary Table 4**. Combined Annotation Dependent Depletion (CADD) score (C-scores) for all SNPs investigated in this study were extracted from n available database [68]. The sheet for all the C-scores is in **Supplementary Table 5**.

### Electrophoretic mobility shift assay (EMSA)

EMSA was performed using the LightShift Chemiluminescent EMSA Kit (Thermo Scientific, catalog number 20148) according to manufacturer instructions. For oligo probe, a 31 bp fragment centered upon the SNP was made by annealing two biotinylated oligonucleotides. Nuclear proteins were extracted from BL2 cells and PBMC using NE-PER Nuclear and Cytoplasmic Extraction Reagents (Thermo Scientific, catalog number 78835) per manufacturer instructions. For gel supershift, 1 μg of Isotype antibody or 10 μl of specific antibodies were added after for additional 30 min incubation. Antibodies used are anti-IKZF1 antibody (Cell Signaling, catalog number 9034S), anti-E2F4 antibody (Cell signaling, catalog number 40291S) and IgG isotype control antibody (Thermo Scientific, catalog number 02-6102).

### Luciferase reporter assay

The luciferase reporter assay was performed exactly according to the manufacturer’s manual (Promega, catalog number E1751). The pGL3 expression vector (0.5 μg) with or without SNPs sequence was co-transfected with the control vector pRL by Nucleofection (Lonza 4D nucleofector) into 5 × 10^5^ Daudi cells. After 48 h incubation, luciferase activity was measured with the Dual-Glo luciferase assay system (Promega, catalog number E2920).

### Pulldown assay and Mass Spectrometry

Pulldown assay is modified from a published protocol for DNA Affinity Purification Assay (DAPA) [26]. Similar to SNP-seq, 500 ng of different alleles (31 bp, biotinylated) from a SNPs were attached to 25 μl streptavidin-Dynabeads (Invitrogen) and then incubated with 100 μg of Daudi nuclear extract for 1 h with or without non-biotinylated competitor (the same alleles). The incubation mixture was then washed in PBST 5 times before eluting with elution buffer (25 mM Tris-Cl PH: 7.5, 1.5M NaCl). The elutions were then precipitated using precipitation kit (EMD Millpore, cat# 539180) and sent for mass spectrometry core at Harvard Medical School. All proteins identified are listed in **Supplementary Table 7**.

### Western blots

Western blots for pulldown assay and knockout validation were performed using corresponding antibodies. Anti-IKZF1 antibody (Cell Signaling, catalog number 9034S), anti-GAPDH antibody (Cell Signaling, catalog number 2118S), anti-IKKε antibody (Cell Signaling, catalog number 2905S), goat anti-Rabbit IgG (H+L) cross-adsorbed secondary antibody, HRP (Thermo Scientific, catalog number G-21234), anti-vinculin antibody (Bio-Rad, catalog number MCA465GA), and goat anti-mouse IgG-HRP (Santa Cruz, catalog number sc-525408).

### Transcriptional factors binding prediction

Candidate TFs binding was analyzed using a published model to identify sequence-specific binding proteins [27, 28].

### Flow cytometry

For IKKε intracellular staining in different subtype cells of PBMCs, cells were treated with or without LPS for four hours and then blocked with Fc Receptor Binding Inhibitor Polyclonal Antibody (eBioscience, catalog number 14-9161-71) and stained with live/dead-e506 stain (eBioscience, 65-0866-14), followed by staining with CD19-APC-e780 (eBioscience, catalog number 47-0199-41); CD14-APC (eBioscience, catalog number 47-0149-41); CD3-BV421 (Biolegend, catalog number 300433); CD4-PE-texas red (eBioscience, catalog number 61-0049-41); CD8-BV786 (eBioscience, catalog number 78-0081-82); CD56-PE-cy7 (eBioscience, catalog number 25-0567-41). anti-IKKε-PE for intracellular staining. All the above antibodies were purchased from eBioscience or labeled manually, used at 1:200 dilution. Purified IKKε antibody was labeled with PE (Biotium, catalog number 92299). For intracellular staining, cells were fixed with Foxp3/Transcription Factor Staining Buffer Set (eBioscience, catalog number 00-5523-00) after surface marker staining. Cells were analyzed on a BD LSRFortessa™ Cell Analyzer (BD Biosciences).

### CRISPR/Cas9 knockout

*IKZF1* in Daudi cells was knocked out using CRISPR-Cas9 through nucleofection (4D nucleofector from Lonza). 1 µl of 40 µM sgRNAs (two sgRNAs to generate a deletion) for *IKZF1* were mixed with 1 µl of 20 µM spCas9 and incubated at 37 degrees for 15 minutes to form ribonucleoprotein (RNP). Formed RNP was kept on ice before use. Daudi cells were then washed with PBS twice and resuspended in SF solution (Lonza, catalog number V4XC-2032), 300k cells per 20 µl solution. Next, RNP was added into each 20 µl solution with Daudi cells and the whole mixture was transferred into nucleofection strip wells for nucleofection with program code CA137. After nucleofection, cells were transferred into pre-warmed culture medium and incubated for 72 hours before analyzing knockout efficiency. sgRNAs used are: 5’-AGGTGGTGGAGAGGTCCTCG-3’ and 5’-TGAGCCCATGCCGATCCCCG-3’.

### CRISPR/Cas9 Homology Directed Repair (HDR)

CRISPR/Cas9 mediated HDR was applied for generation of base-edited cell lines using sgRNA targeting DNA sequences near rs2297550 and asymmetrical single-stranded DNA donors as previously described [69]. 3x10^5^ Daudi cells were nucleofected with 20 picomole of sgRNA-Cas9 complex and 100 picomole of DNA donor template using program CA137 of Amaxa 4D-Nucleofector and SF cell line kit S (Lonza, V4XC-2032). The edited single-cell clones were sorted into 96-well plate by BD Aria II sorter and expanded for 4 weeks. Genotype of individual clones were analyzed by Sanger sequencing to identify C/G wild-type clones (n=3), C/C edited clones (n=4), G/G edited clones (n=3). sgRNA and single-stranded donor DNA for generating CC genotype are 5’-GAGAAAGAGAGAGCGAGAGACGG-3’ and 5’-TTGTGGCGGTGGGACACCAAGCTCAGCACTGAGTCCTGGCTCTCAGCTCTCTCGCACTTCCTGATTTCCGTCTCTCGCTCTCTCTTTCTCTCTTAAAGAGACAGAGGCA-3’; sgRNA and single-stranded donor DNA for generating GG genotype are 5’-TCTTTAAGAGAGAAAGAGAGAGG-3’ and 5’-TTGTGGCGGTGGGACACCAAGCTCAGCACTGAGTCCTGGCTCTCAGCTCTCTCGCACTTCCTGATTTCCGTCTCTCCCTCTCTCTTTCTCTCTTAAAGAGACAGAGGCA-3’.

### RT-qPCR

Total RNA was isolated with Absolutely RNA Miniprep Kit (Agilent, catalog number 400800). cDNA was synthesized with AffinityScript QPCR cDNA Synthesis Kit (Agilent, catalog number 600559). All procedures were performed following the manufacturer’s protocols. RT-qPCR was done with a Agilent AriaMX qPCR machine according to the protocol for Brilliant II SYBR QPCR Low ROX Mstr Mx (Agilent, catalog number 600830). IKZF1 qPCR primers: for k/o clone 1, forward, 5’-CGTGCTGGACAGACTAGCAA - 3’, reverse, 5’-ATTTCGTTCTCCTTCTCGTAGC-3’. IKBKE qPCR primers: forward, 5’-CTTGTGAATCTGCTCCTCGTT-3’, reverse, 5’-GGTACATGAGGACAGAAGCATC-3’.

### Chromatin Immunoprecipitation (ChIP)-qPCR

ChIP was performed for Daudi cells using CHIP-it kit (Active motif, catalog number 53042) according to manufacturer’s protocols. Antibodies used are isotype control antibody and anti-IKZF1 antibody from EMSA. The primers used for Daudi cell CHIP-qPCR to detect rs2297550 are forward, 5’-CTCAGAGCATCCTCTTTCTCTTT -3’, reverse, 5’-CTCTCTCGCACTTCCTGATTT -3’.

### Allele discrimination CHIP-qPCR

CHIP was performed using HDR clones with rs2297550 C/G genotype. Antibodies used are isotype control antibody and anti-IKZF1 antibody from EMSA. Probes and primers used are included in the Custom TaqMan™ SNP Genotyping Assay, human for rs2297550 from ThermoFisher.

### Statistical analyses

All statistical analysis was done using GraphPad software (Prism) using one-way analysis of variance (ANOVA) for comparison of multiple groups, or unpaired t-test (non-parametric Mann-Whitney test) for two groups, with P-values as indicated in the figure legends.

### Data availability

All data supporting the findings of this study are available within the paper and its Supplementary Information. The mass spectrometry proteomics data have been deposited to the ProteomeXchnage Consortium via the PRIDE[70] partner repository (https://www.ebi.ac.uk/pride/) with the dataset identifier PXD048367 and 10.6019/PXD048367.

## Acknowledgements

QW, TK, and M-MB were supported by Joint Biology Consortium microgrants off parent grant NIH/NIAMS P30AR070253. JC was supported by P30AR070253. JAS was supported by NIH/NIAMS R01 AR077607, P30 AR070253, and P30 AR072577, the R. Bruce and Joan M. Mickey Research Scholar Fund, and the Llura Gund Award for Rheumatoid Arthritis Research and Care. MTW was supported by NIH grants R01NS099068, R01HG010730, U01AI130830, R01AI024717, R01AR073228. M-GA was supported by NIH/NIAMS P30AR069625, Gilead Sciences, a Global Team Science Award from the Lupus Research Alliance, and the Arthritis National Research Foundation Vic Braden Family Fellowship. PAN was supported by a Global Team Science Award from the Lupus Research Alliance and NIH/NIAMS 2R01AR065538, R01AR075906, R01AR073201, and P30AR070253. We thank Dr. Kaur Alasoo for making available full summary statistics from the eQTL Catalogue and for recommendations on colocalization analysis.

## Author Contributions

QW and PAN conceived and designed the study. QW, TK, MM-B, SS performed experiments and data analysis. TK led revision experiments. VA, JC, and MG-A performed data analysis. JAS led recruitment of genotyped human subjects. XTC, MTW identified candidate transcription factors. MT, BW performed experiments. QW, TK, PAN drafted the manuscript and all authors edited and approved the manuscript. PAN supervised the research.

## Declaration of interests

The authors declare no relevant competing interests.

## Notes

### Competing Interest Statement

The authors have declared no competing interest.

### Summary of Updates

The list of authors, their order, and affiliations updated; Figure 1 revised.

