## Supplementary Figures for "High-throughput identification of functional regulatory SNPs in systemic lupus erythematosus"

* Co-first authors.

### S. Figure 1

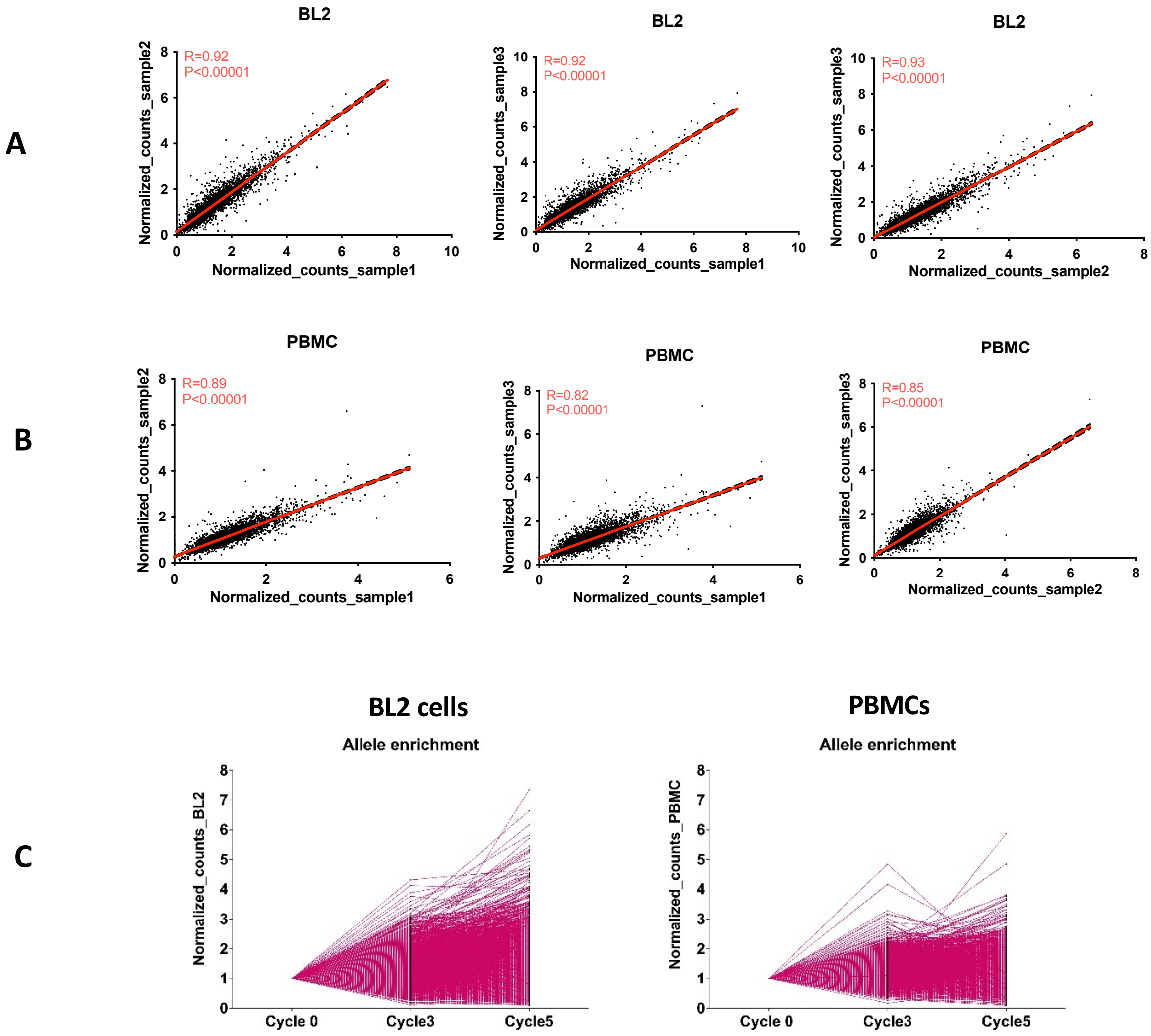

**S Figure 1. Allele-specific enrichment analysis by SNP-seq.** We employed SNP-seq to identify potential regulatory variants from 2180 SNPs utilizing 4363 dsDNA constructs and nuclear extracts from BL2 B cells and PBMCs. **(A, B)** The NGS-generated allele counts normalized to control displayed strong Spearman's correlations among pairs of 3 biological replicates from either BL2 cells or PBMCs. **(C)** The normalized allele1 and allele2 counts ratio for 496 SNPs demonstrated allele-specific enrichment by the protection of nuclear proteins binding across 0-5 cycles.

**S. Figure 2. Analytical approaches to identify SNPs conferring allele-specific protection. (A)** Selection of SNPs demonstrating difference in protection between alleles in cycle 5. **(B)** Selection of SNPs demonstrating an accumulated difference in protection between alleles when comparing cycle 3 and 5.

**A**

cycle 5 of different treatment (Horizontal)

Raw data (read/percentage)

| Cycle 5 | | | | | | | |
|---|---|---|---|---|---|---|---|
| | Mean Percentage in Ctrl | Percentage in BL2 | | | Percentage in PBMC | | |
| | | Replicate 1 | Replicate 2 | Replicate 3 | Replicate 1 | Replicate 2 | Replicate 3 |
| Allele 1 | C1 | B1-1 | B1-2 | B1-3 | P1-1 | P1-2 | P1-3 |
| Allele 2 | C2 | B2-1 | B2-2 | B2-3 | P2-1 | P2-2 | P2-3 |

Sample normalized to control

| Cycle 5 | | | | | | |
|---|---|---|---|---|---|---|
| | Normalized Percentage in BL2 | | | Normailized Percentage in PBMC | | |
| | Replicate 1 | Replicate 2 | Replicate 3 | Replicate 1 | Replicate 2 | Replicate 3 |
| Allele 1 | B1-1/C1 | B1-2/C1 | B1-3/C1 | P1-1/C1 | P1-2/C1 | P1-3/C1 |
| Allele 2 | B2-1/C2 | B2-2/C2 | B2-3/C2 | P2-1/C2 | P2-2/C2 | P2-3/C2 |

Mean Normalized Percentage (MNP)

| Cycle 5 | | |
|---|---|---|
| | Mean Normalized Percentage in BL2 | Mean Normailized Percentage in PBMC |
| Allele 1 | B1-1+B1-2+B1-3/C1/3 | P1-1+P1-2+P1-3/C1/3 |
| Allele 2 | B2-1+B2-2+B2-3/C2/3 | P2-1+P2-2+P2-3/C2/3 |

Remove the alleles have a MNP less than 1 in either BL2 or PBMC

| Total Alleles | Remaining Alleles |
|---|---|
| 4363 | 2053 |

The remaining alleles are divided into two groups:
Group 1: both alleles from one SNPs have a MNP>1;
Group 2: only one allele from one SNPs has a MNP>1.

| Both alleles > 1 | One allele > 1 |
|---|---|
| 1603 | 450 |

Group 1

| Cycle 5 | | |
|---|---|---|
| | BL2 | PBMC |
| MNP ratio | Allele 1/Allele 2 | Allele 1/Allele 2 |

Remove 0.8<A1/A2 ratio <1.2 in either BL2 and PBMC group

| Cycle 5 | |
|---|---|
| Both Alleles >1 | A1/A2>1.2 or A1/A2<0.8 |
| 1603 | 258 (SNPs) |

Group 2
(remove alleles have MNP < 1.2)

| Cycle 5 | |
|---|---|
| Allele >1 | A1 or A2 |
| 450 | 194 (SNPs) |

#### S. Figure 2
(continued)

### B

cycle 3 and 5 of different treatment (Vertical)

Raw data (read/percentage)

↓

| | Mean Percentage in Ctrl | BL2 | | | | | |
|---|---|---|---|---|---|---|---|
| | | Cycle 3 | | | Cycle 5 | | |
| | | Replicate 1 | Replicate 2 | Replicate 3 | Replicate 1 | Replicate 2 | Replicate 3 |
| Allele 1 | C1 | B1.3-1 | B1.3-2 | B1.3-3 | B1.5-1 | P1.5-2 | P1.5-3 |
| Allele 2 | C2 | B2.3-1 | B2.3-2 | B2.3-3 | P2.5-1 | P2.5-2 | P2.5-3 |

↓

Sample normalized to control, remove the alleles have MNP<1

↓

| Total Alleles | Remaining Alleles |
|---|---|
| 4363 | 1903 |

↓

The remaining alleles are divided into two groups:
Group 1: both alleles from one SNPs have a MNP>1;
Group 2: only one allele from one SNPs has a MNP>1.

↓

| Both alleles > 1 | One allele > 1 |
|---|---|
| 1474 | 429 |

Group 1

| BL2 | | |
|---|---|---|
| | Cycle 3 | Cycle 5 |
| MNP ratio | Allele 1/Allele 2 | Allele 1/Allele 2 |

↓ Keep the alleles has an increasing or decreasing ratio of A1/A2 from cycle 3 to cycle5

| Both alleles > 1 | Increasing or Decreasing MNP ratio |
|---|---|
| 1474 | 297 (SNPs) |

Group 2
(Keep the alleles has increasing MNP)

| BL2 | |
|---|---|
| Allele >1 | A1 or A2 |
| 450 | 199 (SNPs) |

Summary

| | Allele Specific Protection | |
|---|---|---|
| | Group 1 | Group 2 |
| Horizontal | 258 | 194 |
| Vertical | 297 | 199 |
| Overlapping | 152 | 96 |

**S. Figure 3**

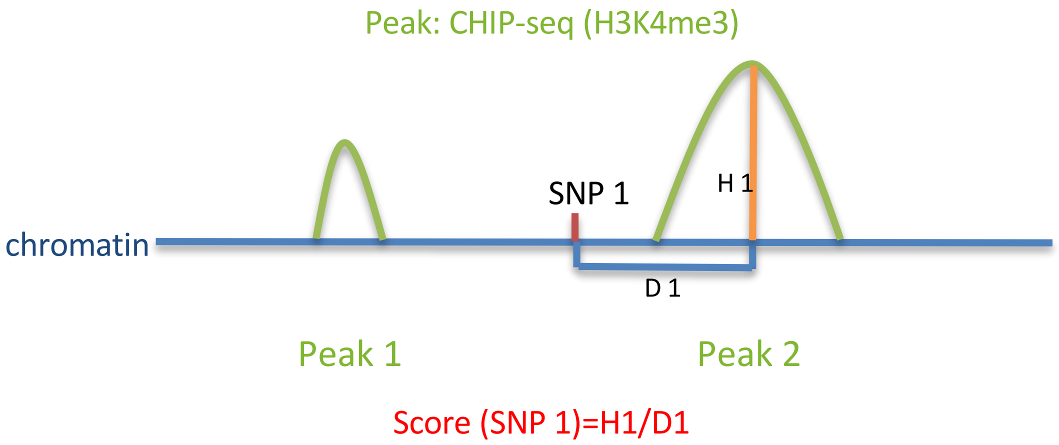

**S. Figure 3. Calculation of H3K4me3 chromatin scores from 5 ChIP-seq studies in human B cells.** SNP locus were scored based on the height of the closest H3K4me3 peak divided by the distance from the SNP to that peak, using 5 studies from the Encyclopedia of DNA Elements (ENCODE) database:

experiment 1 https://www.encodeproject.org/experiments/ENCSR878JSF/
experiment 2 https://www.encodeproject.org/experiments/ENCSR939UQD/
experiment 3 https://www.encodeproject.org/experiments/ENCSR269OVV/
experiment 4 https://www.encodeproject.org/experiments/ENCSR000DQR/
experiment 5 https://www.encodeproject.org/experiments/ENCSR000DQP/

**S. Figure 4. electrophoretic mobility shift assay (EMSA) blots for 52 candidate SNPs.** Biotinylated 31 bp DNA probes centered on each allele of the 52 SNPs were incubated with nuclear extracts from either BL2 B cells or PBMCs for the EMSA. Non-biotinylated DNA fragment served as competitor.

Please see plots at end of Supplemental Figures.

**S. Figure 5. Summary table of EMSA validation experiments for 52 candidate SNPs.** 52 SNPs were initially assessed through EMSA using BL2 nuclear extract. Among them, 21 SNPs showed consistent allelic differential binding across both technical and biological replicates. 9 SNPs were further confirmed using nuclear extract from PBMCs**.** Lead SNPs identified as SLE GWAS hits were marked in red. NE; nuclear extract. Please see Suppl. Fig. 4 for plots.

| | Allele specific binding | | | Allele specific binding | Allele specific binding |
|---|---|---|---|---|---|
| SNPs | BL2 NE from Sept. | | BL2 NE from Oct. | (summary for BL2) | PBMC NE from Sept. |
| | Exp. 1 | Exp.2 | Exp.3 | | |
| rs2297550 | √ | √ | √ | √ | √ |
| rs56741490 | √ | √ | √ | √ | |
| rs61823882 | √ | √ | × | √ | |
| rs4952115 | × | × | × | × | |
| rs906868 | √ | √ | √ | √ | √ |
| rs1132200 | √ | √ | √ | √ | |
| rs157042 | × | × | × | × | |
| rs7769961 | × | √ | √ | √ | |
| rs13205210 | × | × | × | × | |
| rs6934662 | × | × | × | × | |
| rs13215181 | × | × | × | × | |
| rs10761604 | × | × | × | × | |
| rs28364617 | × | × | × | × | |
| rs11227302 | × | × | × | × | |
| rs11606631 | × | × | × | × | |
| rs1728769 | √ | √ | × | × | |
| rs1728770 | × | × | × | × | |
| rs58363746 | √ | √ | √ | √ | |
| rs936394 | √ | √ | √ | √ | √ |
| rs113370572 | √ | × | × | × | |
| rs180769054 | √ | √ | × | × | |
| rs118126443 | × | × | × | × | |
| rs11089620 | × | × | × | × | |
| rs2298428 | × | × | × | × | |
| Red: Lead SNPs | | | | | |

| | Allele specific binding | | | Allele specific binding | Allele specific binding |
|---|---|---|---|---|---|
| SNPs | BL2 NE from Oct. | | BL2 NE from Sept. | (summary for BL2) | PBMC NE from Sept. |
| | Exp. 1 | Exp.2 | Exp.3 | | |
| rs17321999 | × | √ | × | × | |
| rs1040029 | √ | √ | √ | √ | |
| rs528765 | √ | √ | √ | √ | |
| rs13360613 | √ | √ | √ | √ | |
| rs276461 | √ | √ | √ | √ | √ |
| rs13213604 | √ | √ | √ | √ | √ |
| rs1645935 | × | × | × | × | |
| rs6501784 | √ | √ | × | × | |
| rs9894009 | × | × | × | × | |
| rs4660115 | √ | √ | √ | √ | |
| rs3768056 | √ | √ | √ | √ | √ |
| rs574808 | × | × | × | × | |
| rs6937876 | √ | √ | √ | √ | |
| rs2814955 | √ | √ | √ | √ | √ |
| rs7745097 | × | × | × | × | |
| rs205286 | × | × | × | × | |
| rs9469877 | × | × | × | × | |
| rs7302634 | √ | √ | √ | √ | √ |
| rs56928975 | × | × | × | × | |
| rs9913957 | √ | √ | √ | √ | |
| rs73304123 | √ | × | × | × | |
| rs9907966 | √ | √ | √ | √ | √ |
| rs9907564 | √ | √ | √ | √ | |
| rs112561568 | × | × | × | × | |
| rs200906779 | × | × | × | × | |
| rs61134033 | √ | × | × | × | |
| rs131664 | × | × | × | × | |
| rs5754467 | × | × | × | × | |
| Red: Lead SNPs | | | | | |

**S. Figure 6. Pulldown and silver staining of 8 EMSA-validated SNPs.** We performed pulldown assays using 31bp oligonucleotides for each allele of the 8 SNPs using nuclear extract from Daudi cells, with or without an excess of non-biotinylated competitor. Silver staining was employed to visualize the allele-specific binding of eluted proteins that were separated by gel electrophoresis. In two biological replicates, four variants showing differential binding were indicated with red circles.

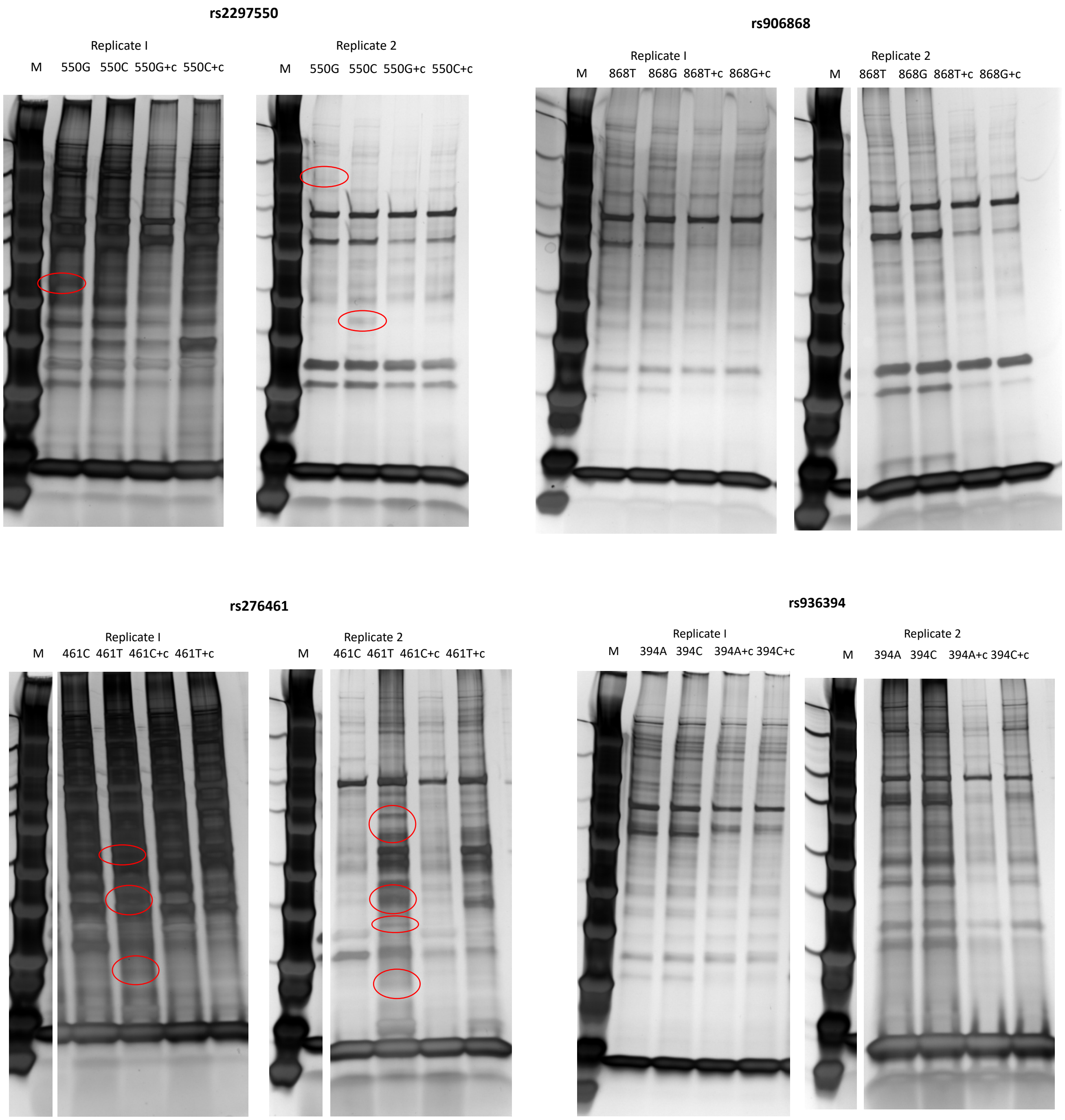

### S. Figure 6 (continued)

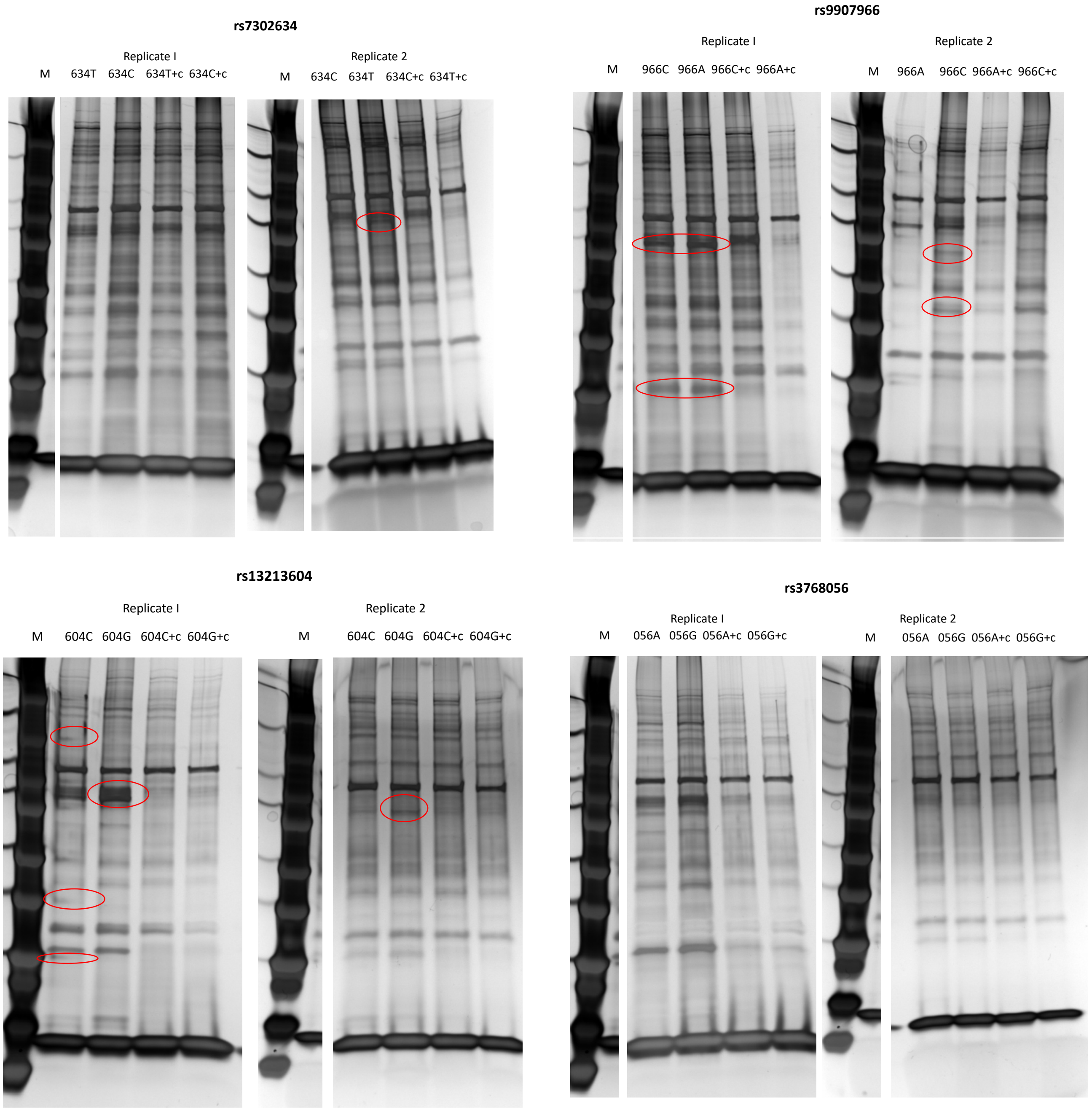

#### S. Figure 7

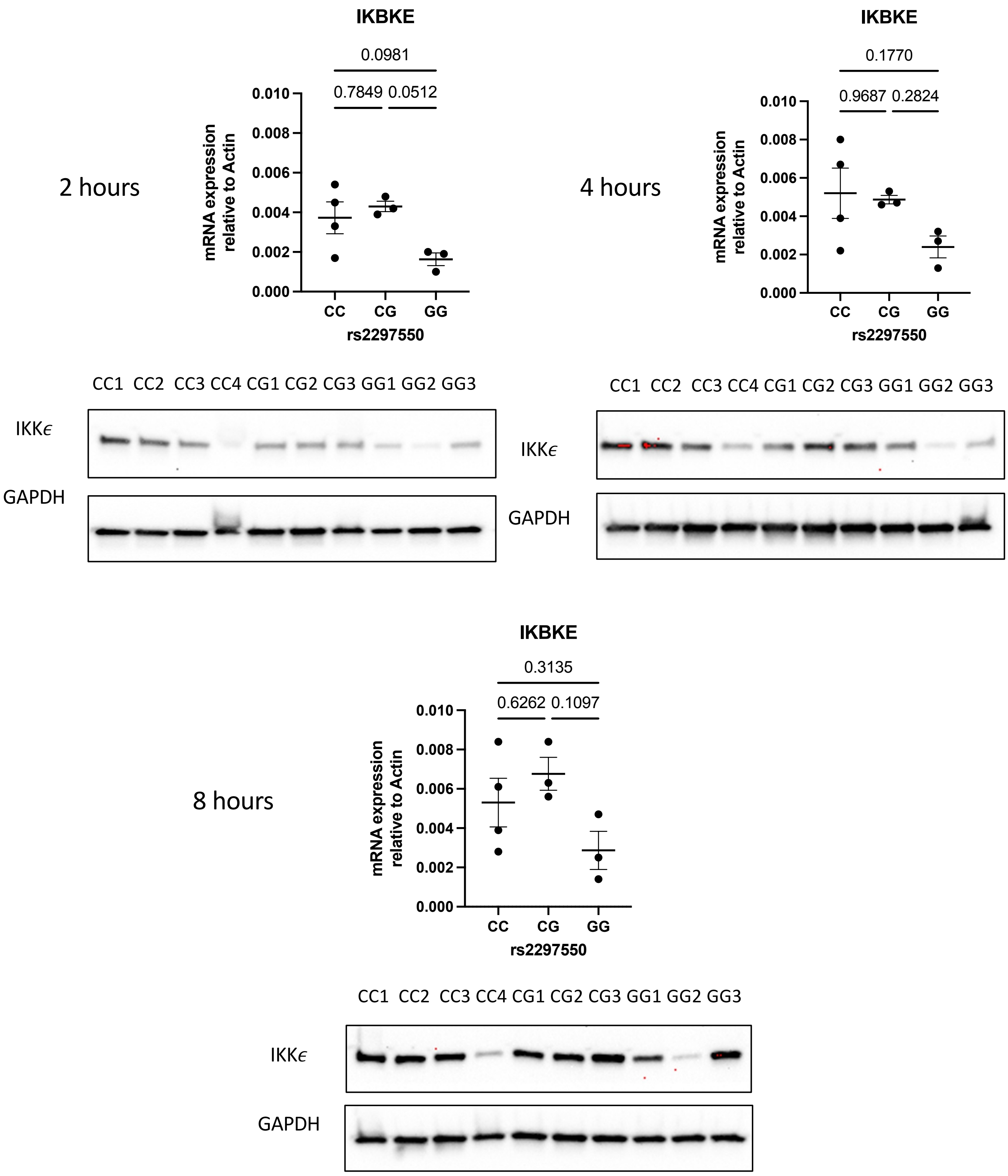

**S. Figure 7. Relationship between genotype and IKKε expression in LPS-stimulated Daudi cells.** Base-edited Daudi cells from Figure 4 were stimulated with LPS 5 $\mu$g/mL for the intervals shown to determine RNA for *IKBKE* and Western blot for IKKε protein.

#### S. Figure 8

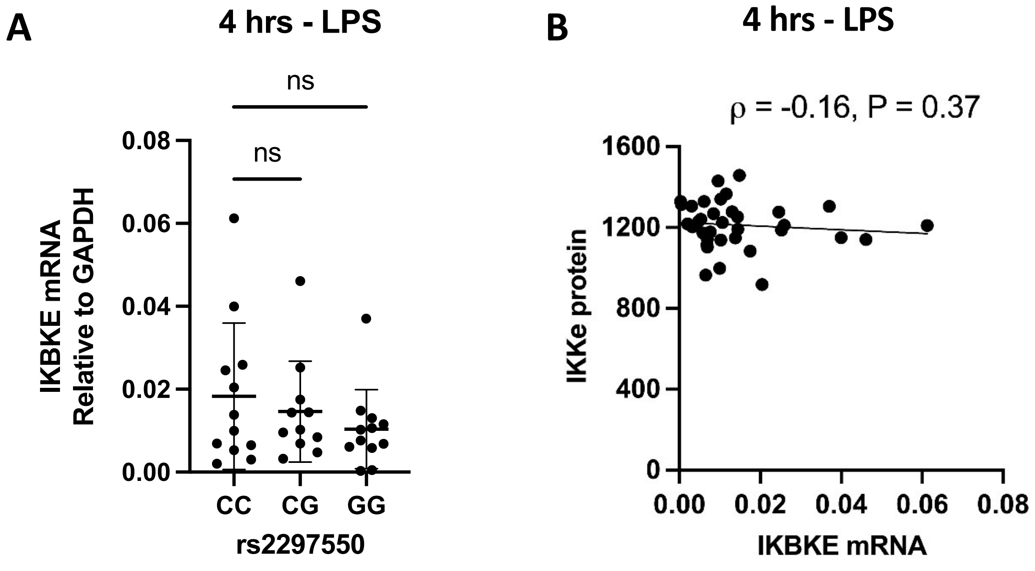

**S. Figure 8. Genotype at rs2297550 does not correlates with *IKBKE* mRNA expression in PBMCs stimulated with LPS for 4 hours. (A)** RNA level of *IKBKE* was determined by RT-qPCR for 35 healthy subjects with C/C (n=12), C/G (n=11), G/G (n=12) genotype at SNP rs2297550. **(B)** Correlation between IKKe protein level (mean fluorescence intensity) and *IKBKE* RNA level (normalized to GAPDH) was determined by Spearman's rank correlation with a two-tailed test.

#### S. Figure 9

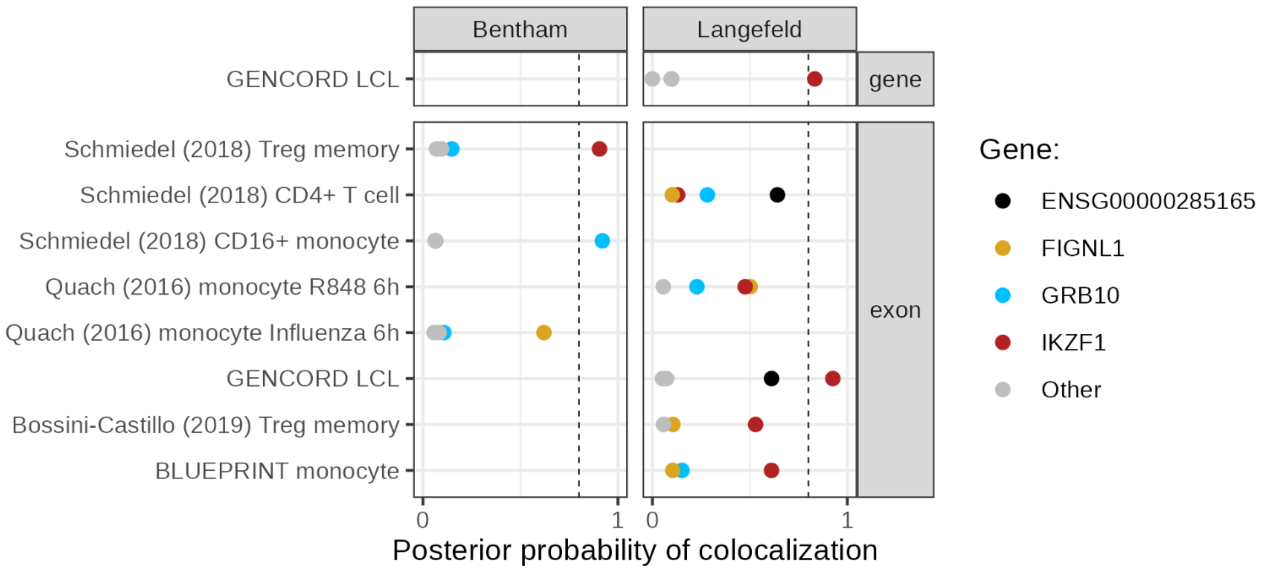

**S. Figure 9. Colocalization between SLE GWAS data and QTLs from the eQTL Catalogue at the *IKZF1* locus.** We selected eQTL data on immune cell types from eQTL Catalogue release 6[1], including molecular traits (genes or exons) within 500kb from the risk variant and with at least one QTL variant passing 5% FDR. We obtained GWAS summary statistics from Bentham et al[2] and Langefeld et al[3]. We tested for colocalization with coloc v5.2.3[4]. The x-axis shows posterior probability of hypothesis 4 (PP4) from coloc. These colocalizations would not be observed if more stringent significance thresholds were used, for example, $p < 1x10^{-5}$ for eQTLs and gene-level FDR for exon-QTLs to account for the high number of molecular traits tested per gene, therefore they should be considered only suggestive evidence for colocalization.

1. Kerimov, N. *et al.* A compendium of uniformly processed human gene expression and splicing quantitative trait loci. *Nat. Genet.* **53**, 1290–1299 (2021).
2. Bentham, J. *et al.* Genetic association analyses implicate aberrant regulation of innate and adaptive immunity genes in the pathogenesis of systemic lupus erythematosus. *Nat. Genet.* **47**, 1457–1464 (2015).
3. Langefeld, C. D. *et al.* Transancestral mapping and genetic load in systemic lupus erythematosus. *Nat. Commun.* **8**, 16021 (2017).
4. Giambartolomei, C. *et al.* Bayesian test for colocalisation between pairs of genetic association studies using summary statistics. *PLoS Genet.* **10**, e1004383 (2014).

#### S. Figure 4

##### EMSA using BL2 cell NE

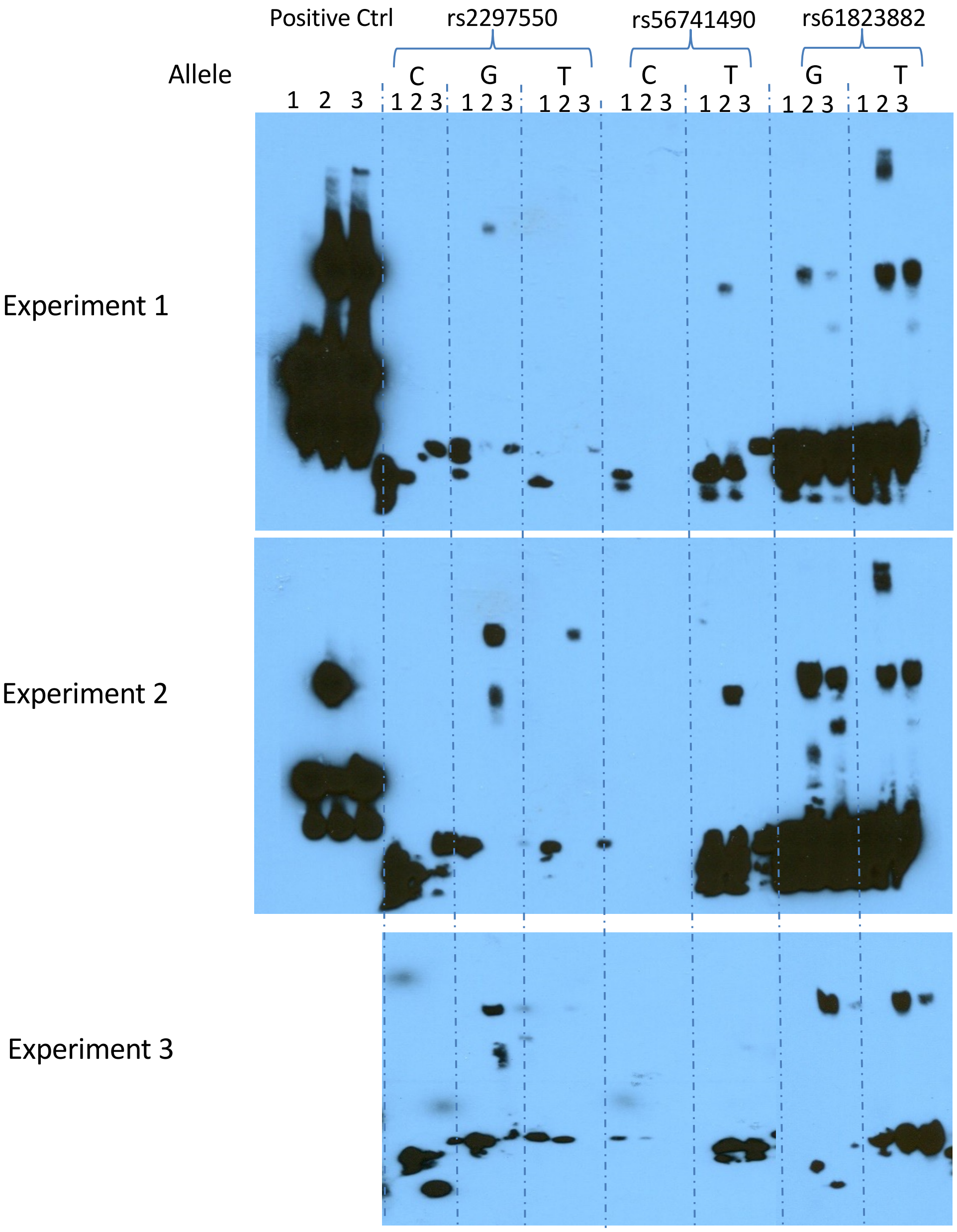

Lane 1: Biotinylated DNA
Lane 2: Biotinylated DNA + Nuclear extract
Lane 3: Biotinylated DNA + Nuclear extract + Non-biotinylated competitor

#### EMSA using BL2 cell NE

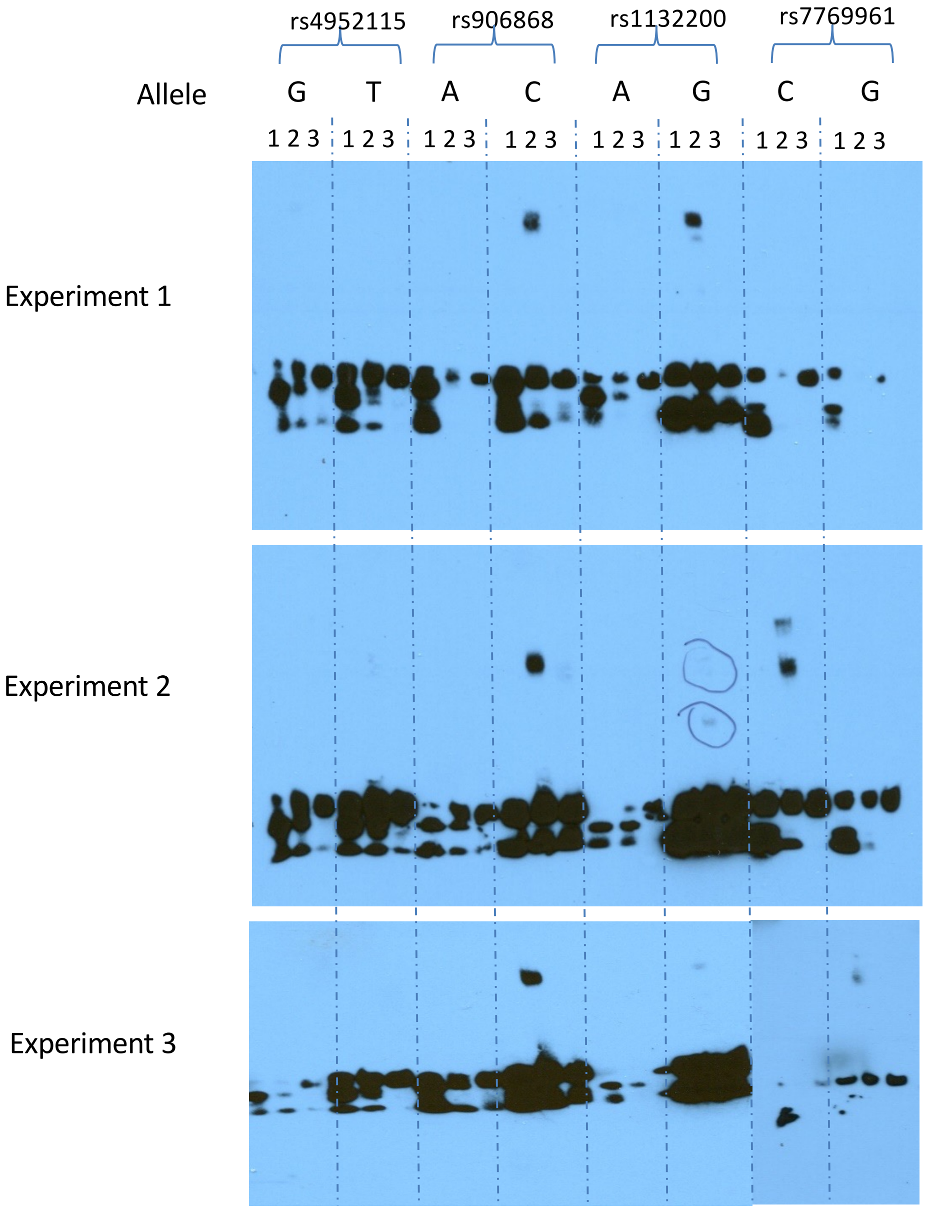

Lane 1: Biotinylated DNA
Lane 2: Biotinylated DNA + Nuclear extract
Lane 3: Biotinylated DNA + Nuclear extract + Non-biotinylated competitor

### EMSA using BL2 cell NE

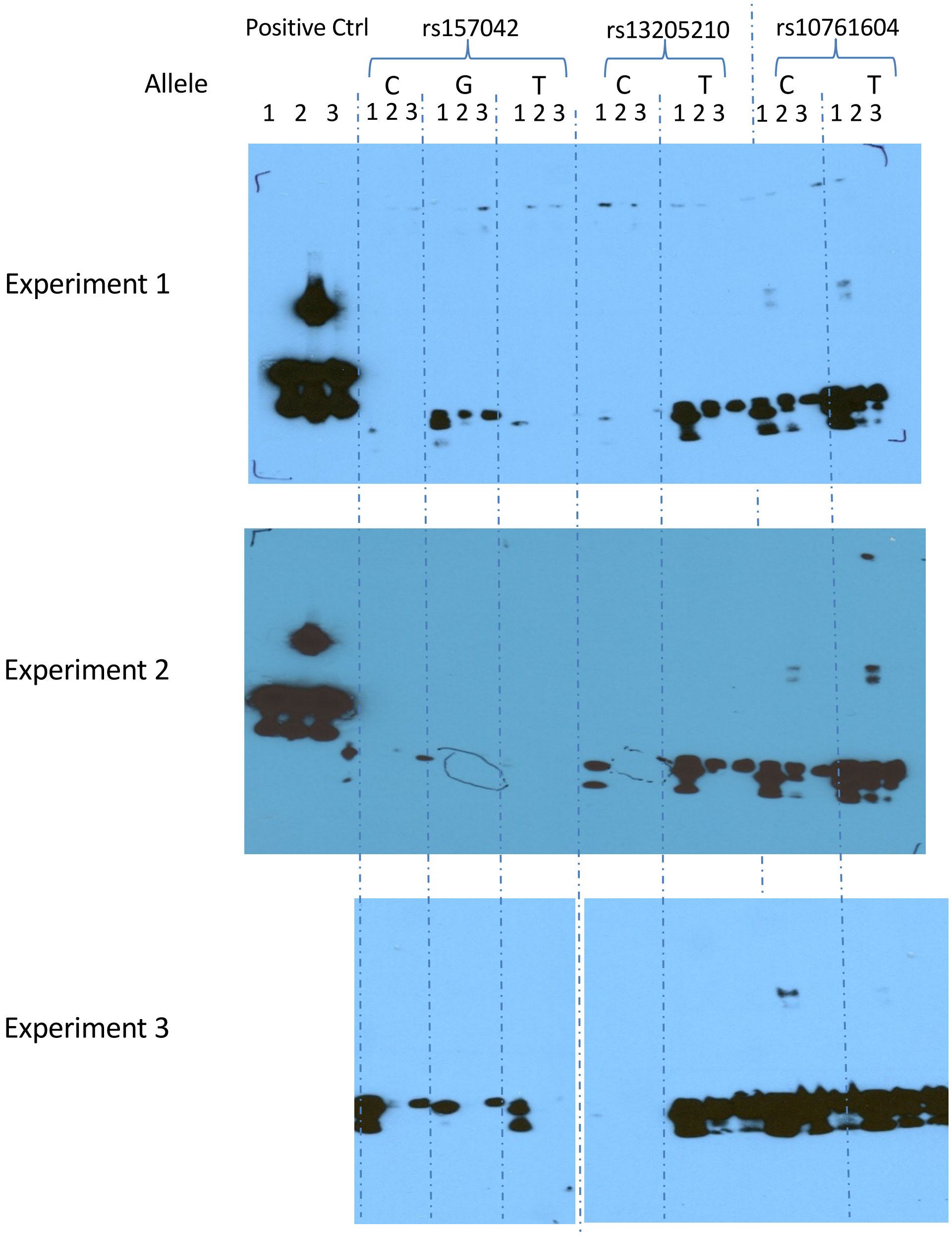

Lane 1: Biotinylated DNA
Lane 2: Biotinylated DNA + Nuclear extract
Lane 3: Biotinylated DNA + Nuclear extract + Non-biotinylated competitor

### EMSA using BL2 cell NE

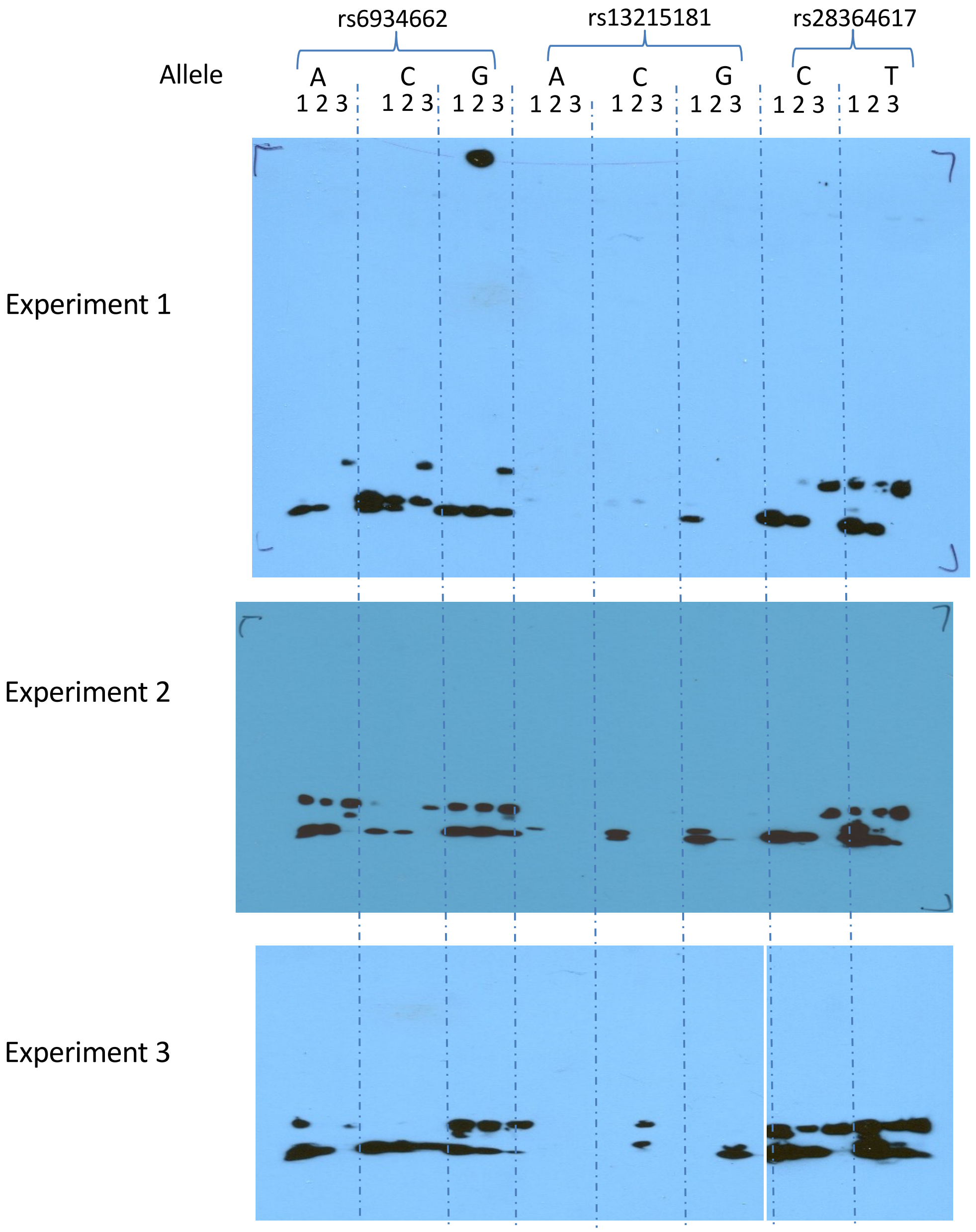

Lane 1: Biotinylated DNA
Lane 2: Biotinylated DNA + Nuclear extract
Lane 3: Biotinylated DNA + Nuclear extract + Non-biotinylated competitor

### EMSA using BL2 cell NE

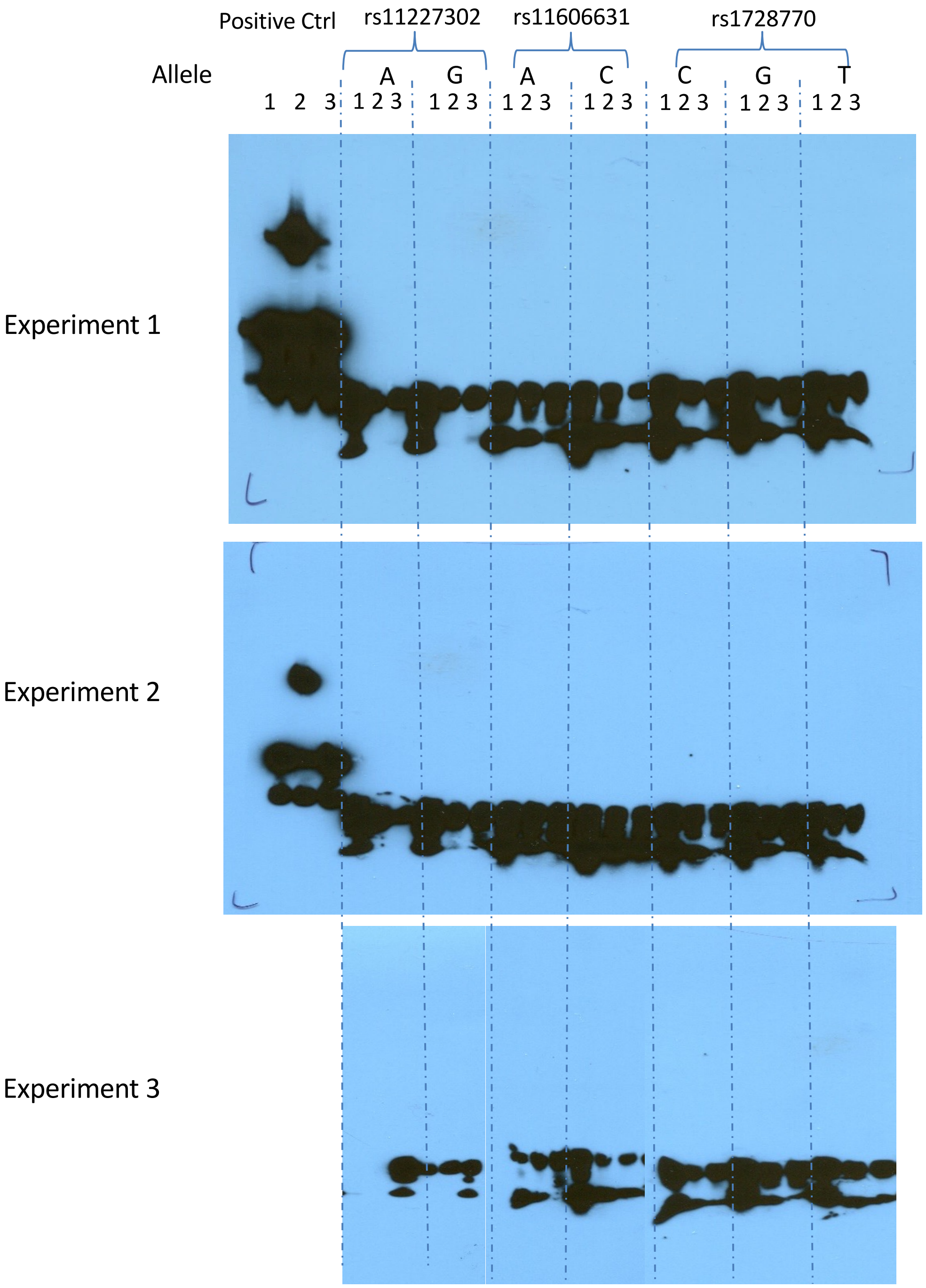

Lane 1: Biotinylated DNA
Lane 2: Biotinylated DNA + Nuclear extract
Lane 3: Biotinylated DNA + Nuclear extract + Non-biotinylated competitor

### EMSA using BL2 cell NE

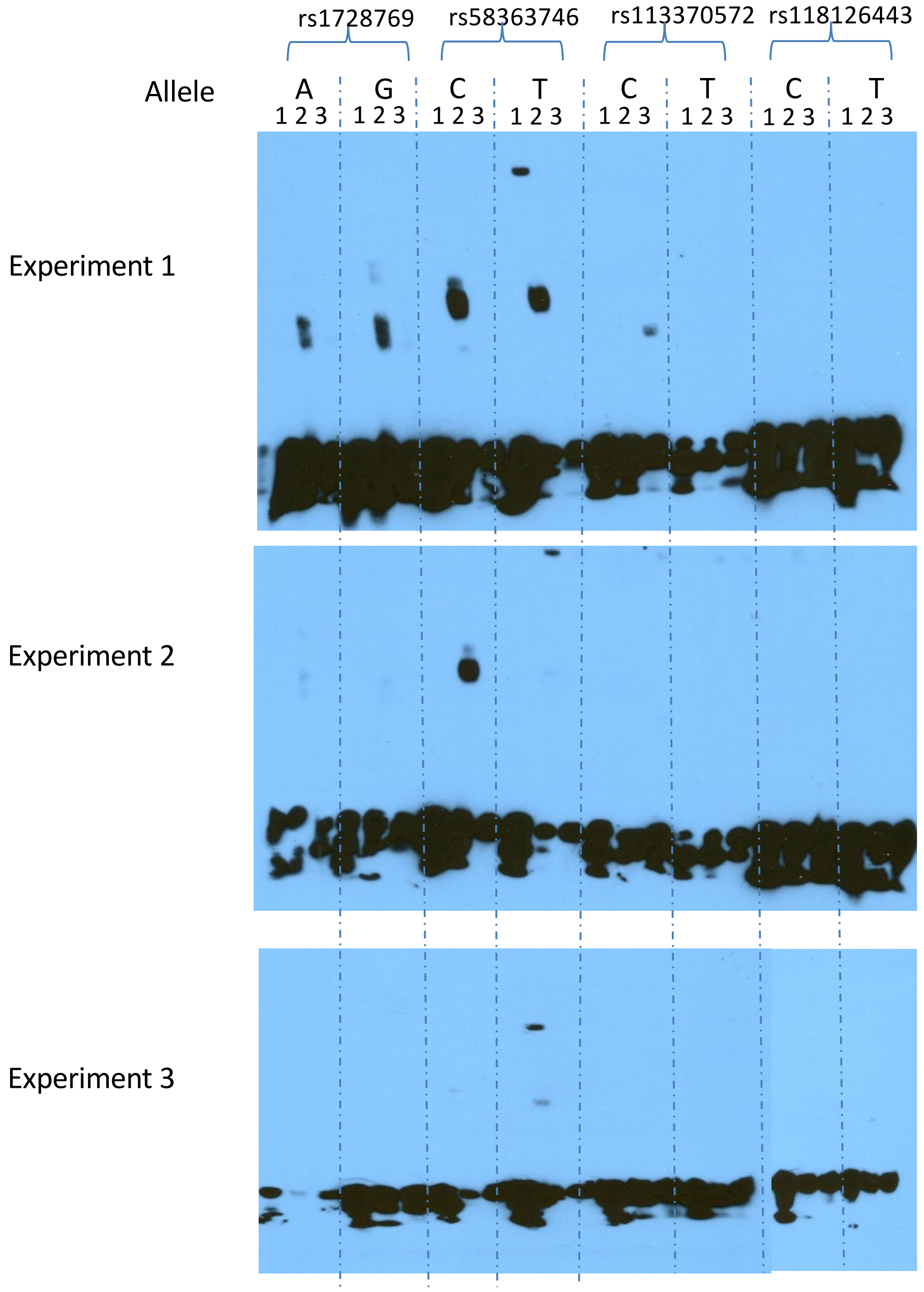

Lane 1: Biotinylated DNA
Lane 2: Biotinylated DNA + Nuclear extract
Lane 3: Biotinylated DNA + Nuclear extract + Non-biotinylated competitor

### EMSA using BL2 cell NE

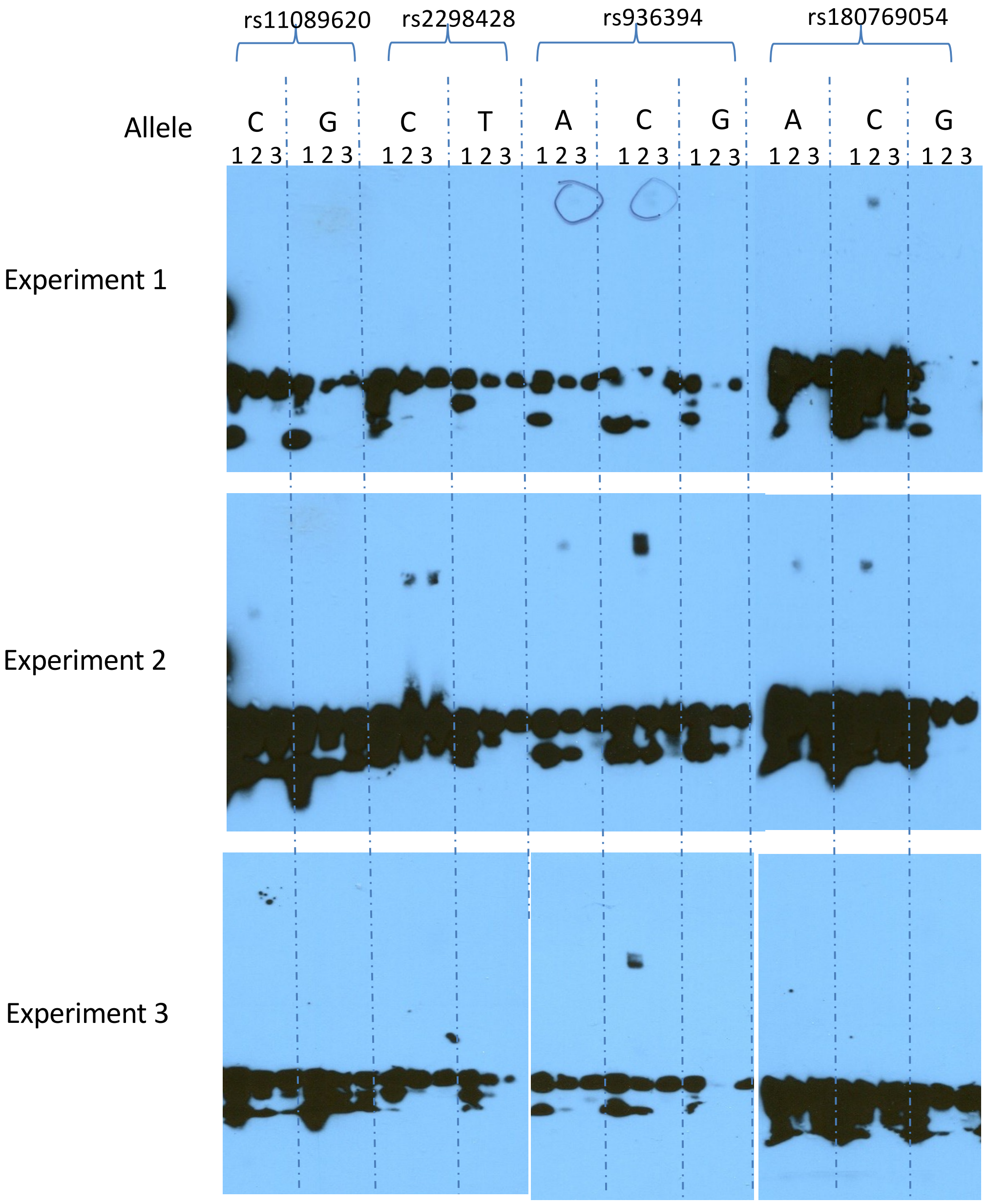

Lane 1: Biotinylated DNA
Lane 2: Biotinylated DNA + Nuclear extract
Lane 3: Biotinylated DNA + Nuclear extract + Non-biotinylated competitor

### EMSA using BL2 cell NE

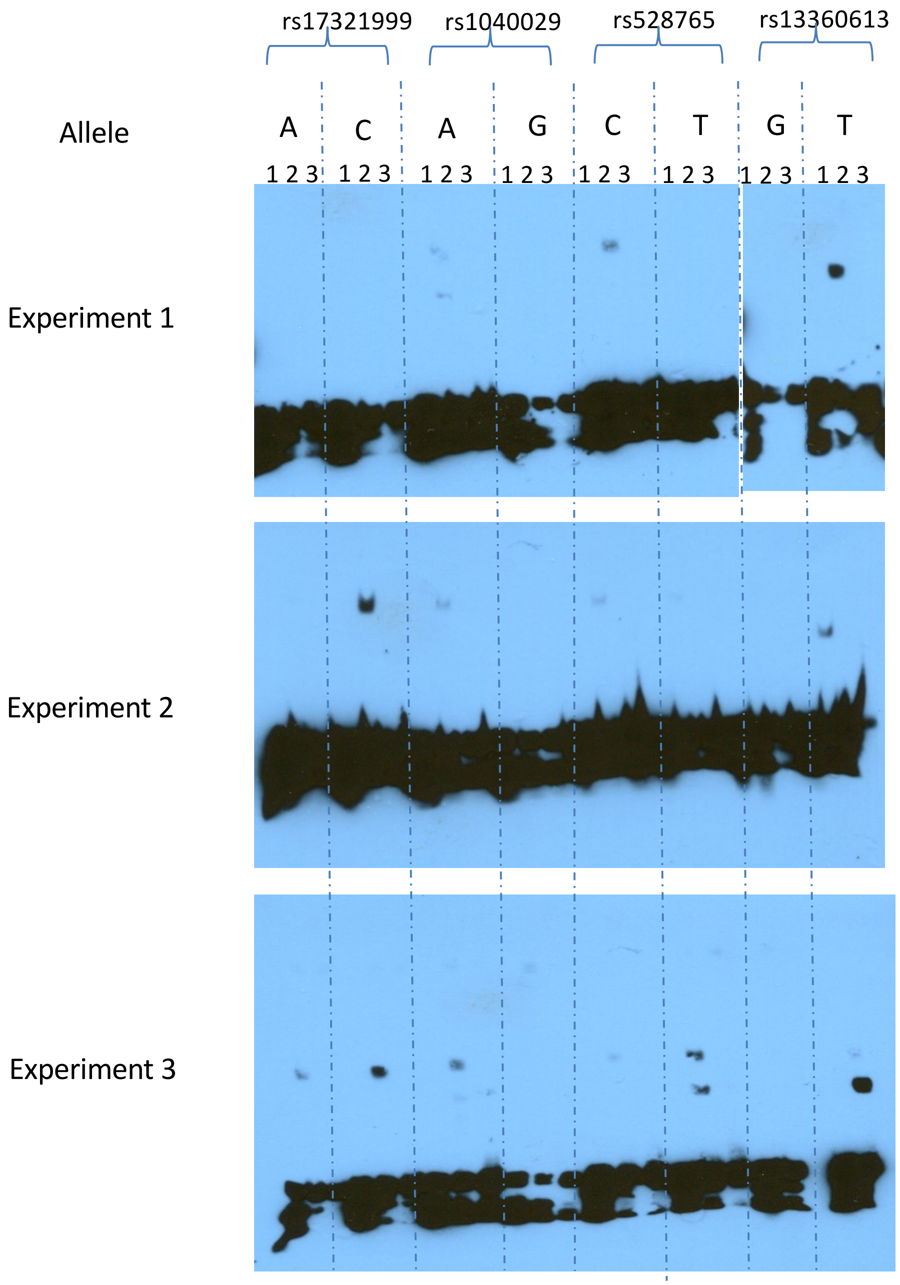

Lane 1: Biotinylated DNA
Lane 2: Biotinylated DNA + Nuclear extract
Lane 3: Biotinylated DNA + Nuclear extract + Non-biotinylated competitor

### EMSA using BL2 cell NE

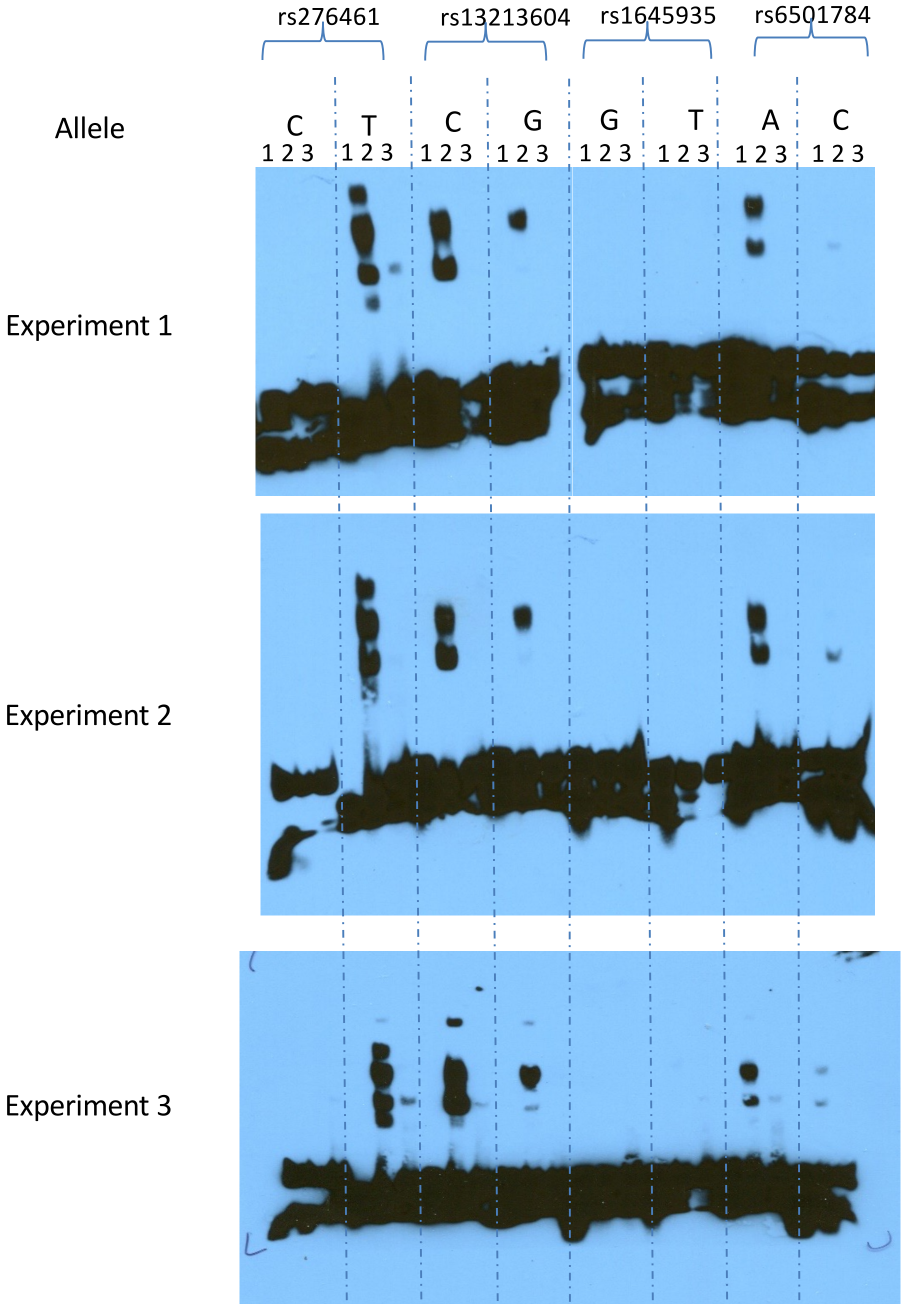

Lane 1: Biotinylated DNA
Lane 2: Biotinylated DNA + Nuclear extract
Lane 3: Biotinylated DNA + Nuclear extract + Non-biotinylated competitor

#### EMSA using BL2 cell NE

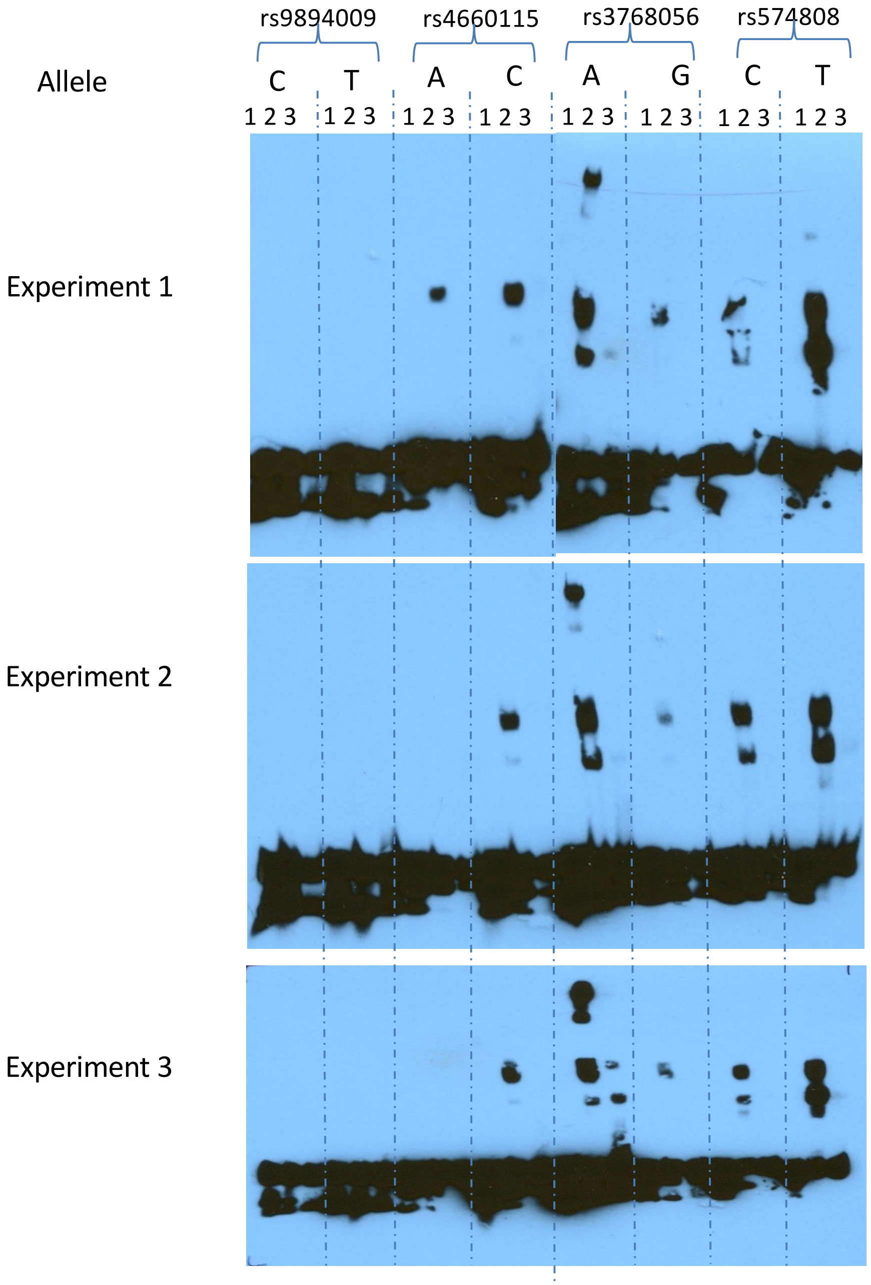

Lane 1: Biotinylated DNA
Lane 2: Biotinylated DNA + Nuclear extract
Lane 3: Biotinylated DNA + Nuclear extract + Non-biotinylated competitor

### EMSA using BL2 cell NE

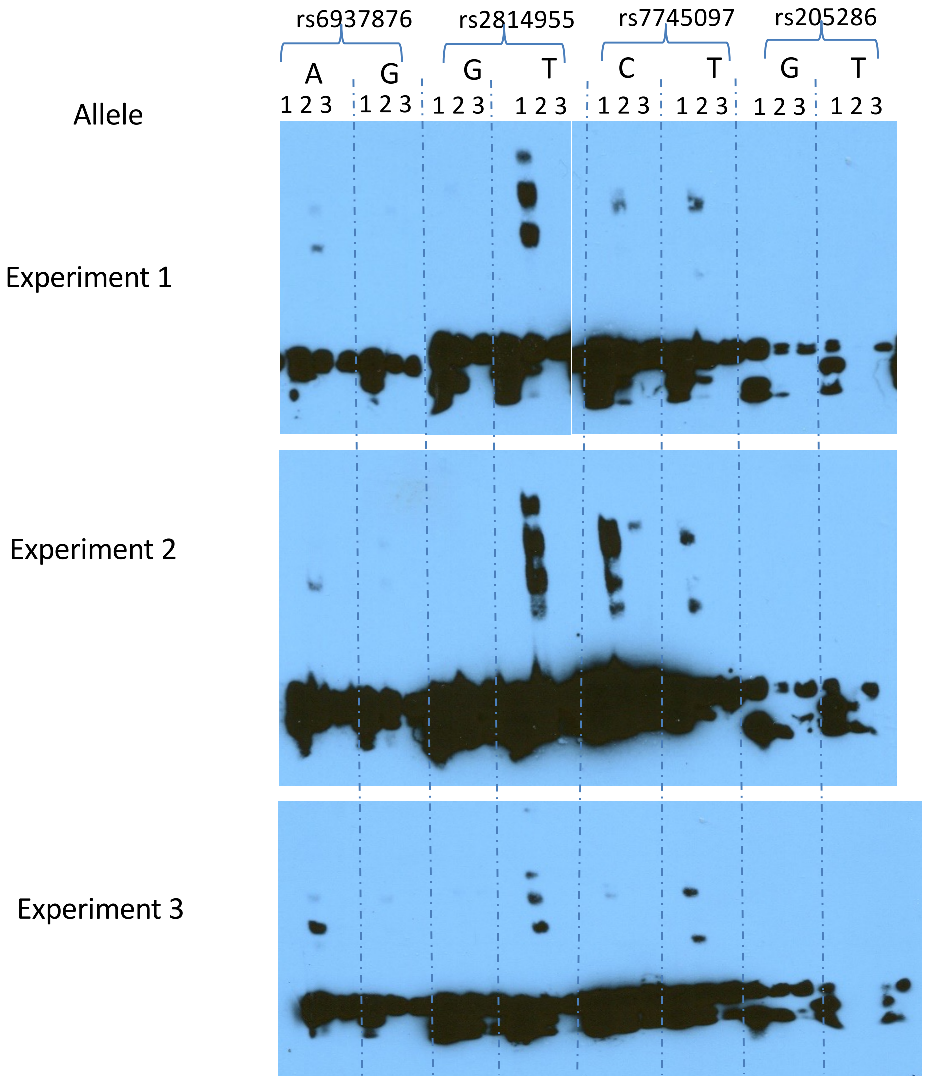

Lane 1: Biotinylated DNA
Lane 2: Biotinylated DNA + Nuclear extract
Lane 3: Biotinylated DNA + Nuclear extract + Non-biotinylated competitor

#### EMSA using BL2 cell NE

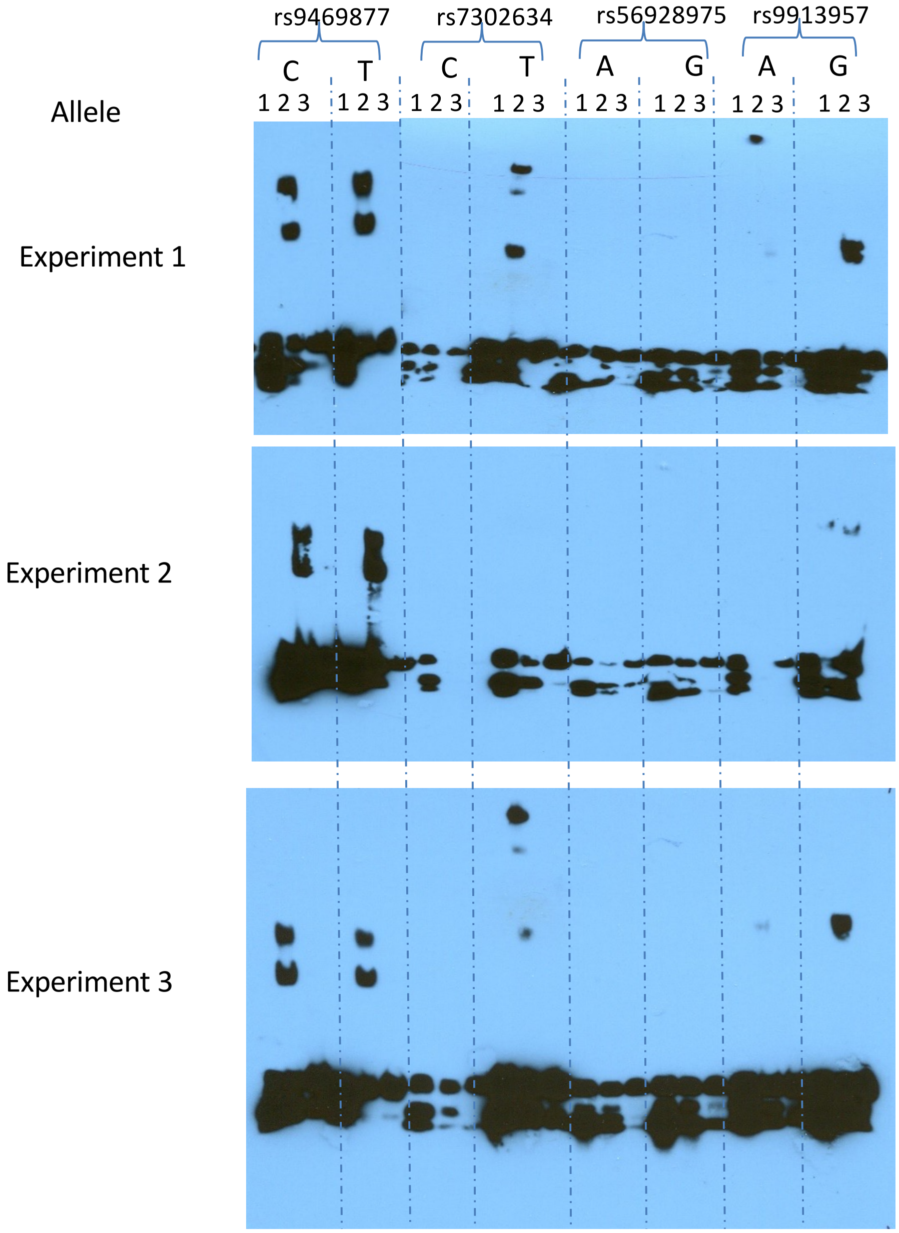

Lane 1: Biotinylated DNA
Lane 2: Biotinylated DNA + Nuclear extract
Lane 3: Biotinylated DNA + Nuclear extract + Non-biotinylated competitor

#### EMSA using BL2 cell NE

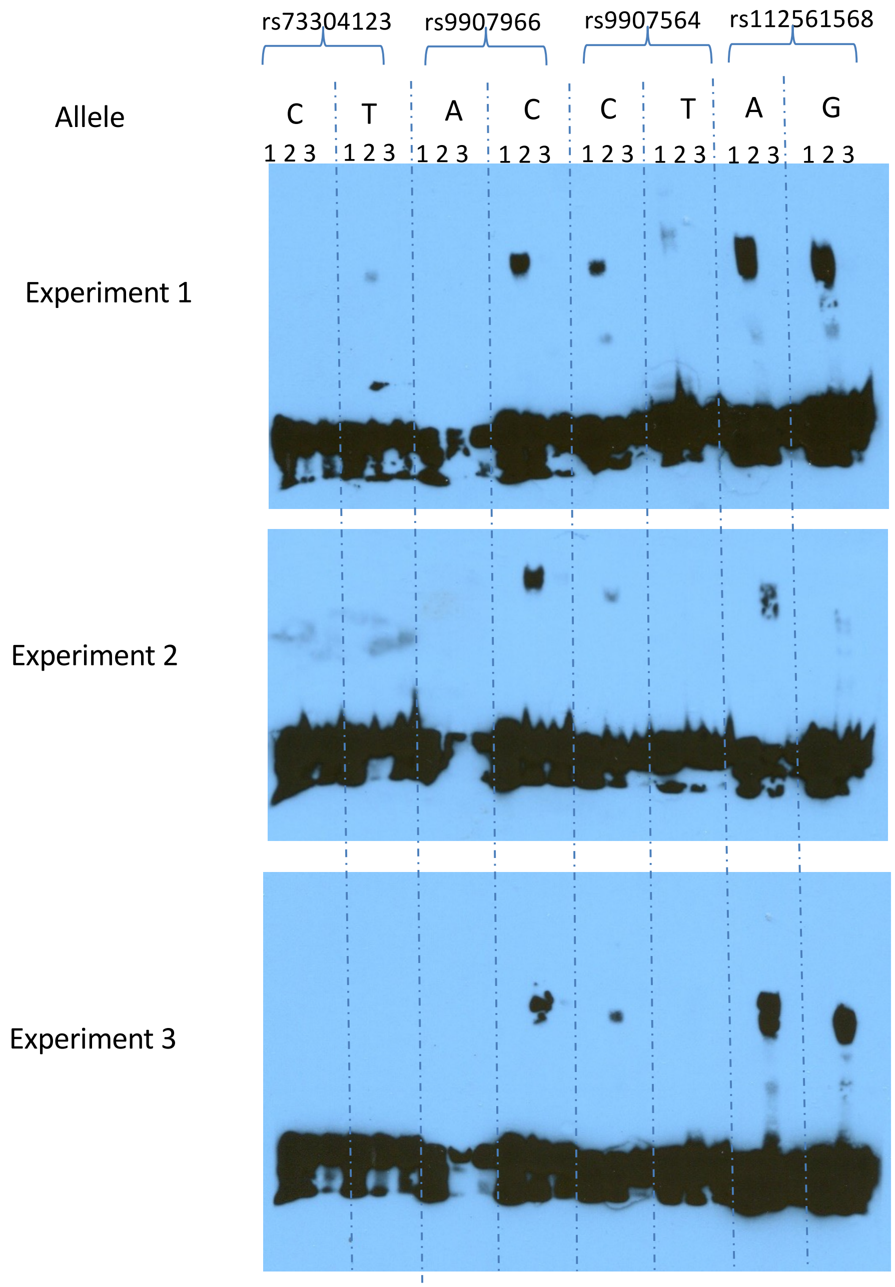

Lane 1: Biotinylated DNA
Lane 2: Biotinylated DNA + Nuclear extract
Lane 3: Biotinylated DNA + Nuclear extract + Non-biotinylated competitor

### EMSA using BL2 cell NE

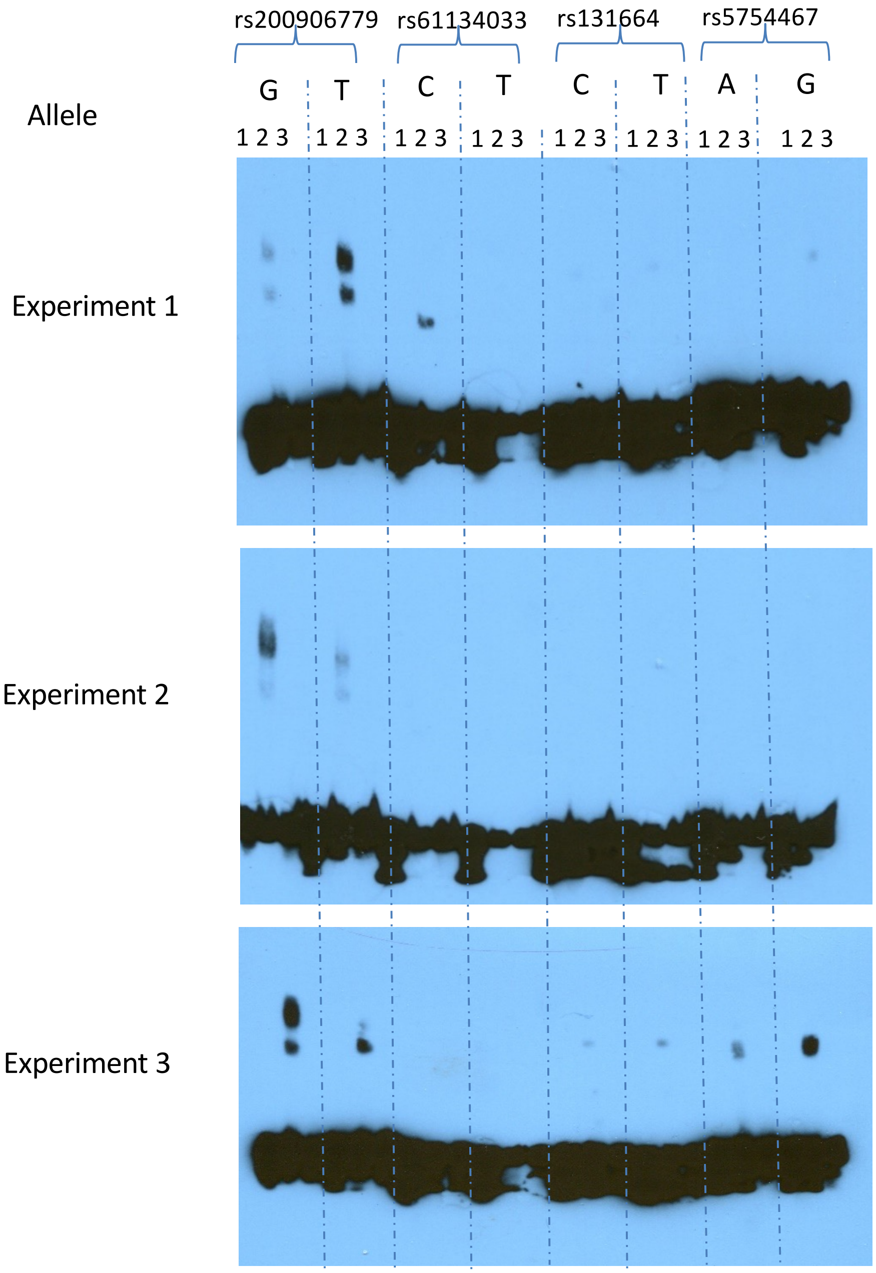

Lane 1: Biotinylated DNA
Lane 2: Biotinylated DNA + Nuclear extract
Lane 3: Biotinylated DNA + Nuclear extract + Non-biotinylated competitor

### EMSA using PBMC NE

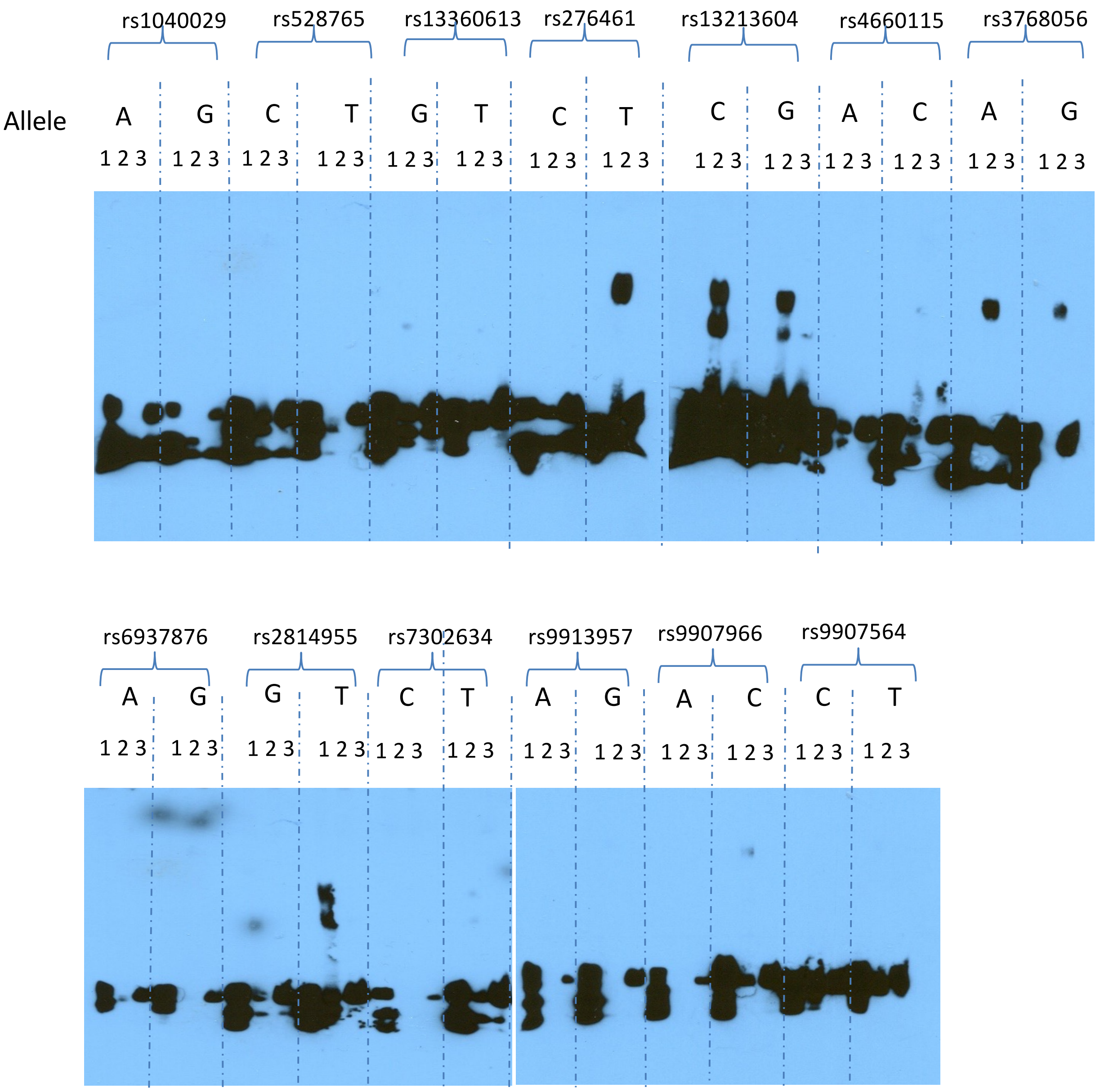

Lane 1: Biotinylated DNA
Lane 2: Biotinylated DNA + Nuclear extract
Lane 3: Biotinylated DNA + Nuclear extract + Non-biotinylated competitor

### EMSA using PBMC NE

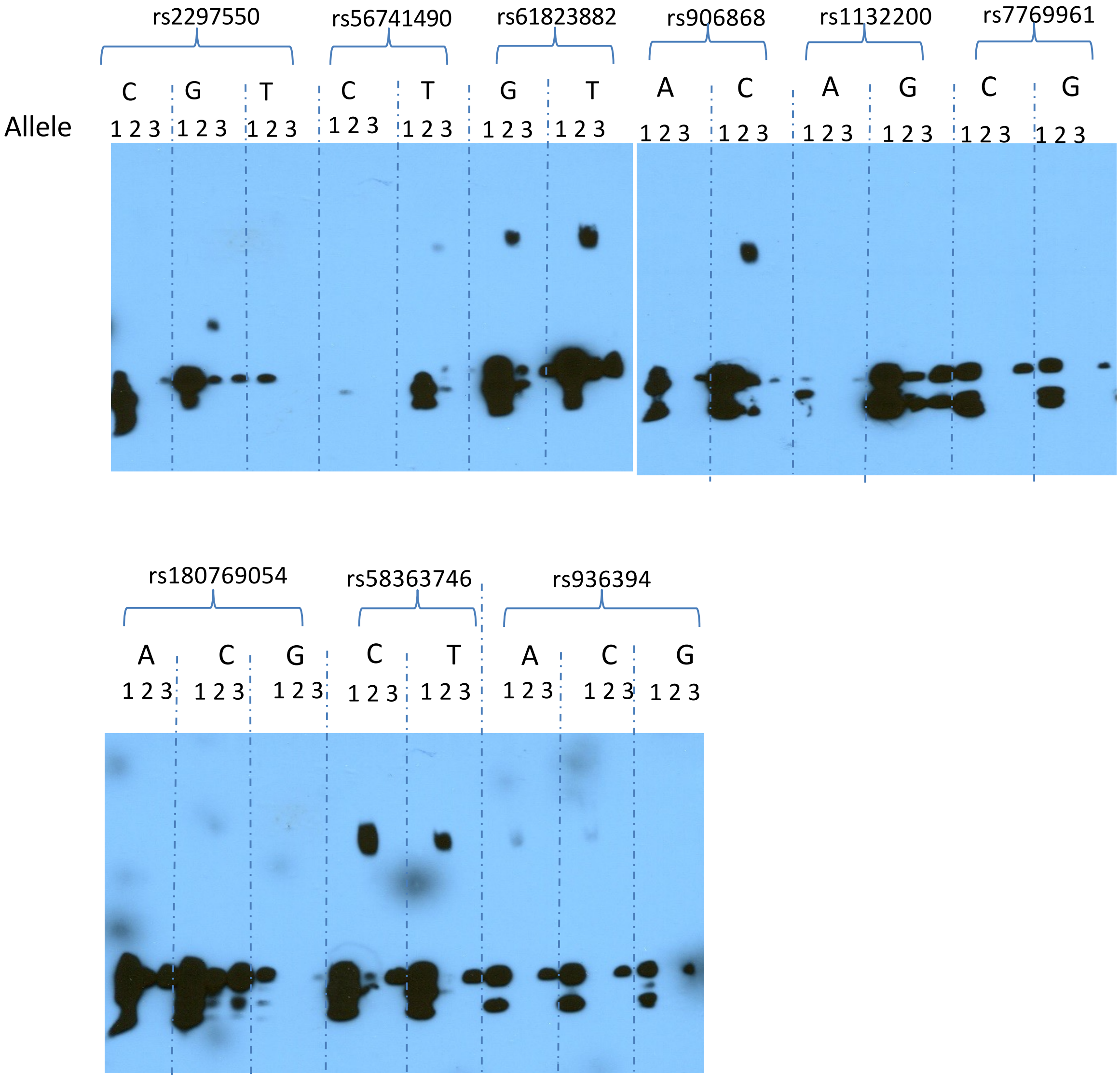

Lane 1: Biotinylated DNA
Lane 2: Biotinylated DNA + Nuclear extract
Lane 3: Biotinylated DNA + Nuclear extract + Non-biotinylated competitor
